# Identification of BAHD acyltransferases associated with acylinositol biosynthesis in *Solanum quitoense* (naranjilla)

**DOI:** 10.1101/2022.03.21.485185

**Authors:** Bryan J. Leong, Steven Hurney, Paul Fiesel, Thilani M. Anthony, Gaurav Moghe, A. Daniel Jones, Robert L. Last

## Abstract

Plants make a variety of specialized metabolites that can mediate interactions with animals, microbes and competitor plants. Understanding how plants synthesize these compounds enables studies of their biological roles by manipulating their synthesis *in vivo* as well as producing them *in vitro*. Acylsugars are a group of protective metabolites that accumulate in the trichomes of many Solanaceae family plants. Acylinositol biosynthesis is of interest because it appears to be restricted to a subgroup of species within the Solanum genus. Previous work characterized a triacylinositol acetyltransferase involved in acylinositol biosynthesis in the Andean fruit plant *Solanum quitoense* (lulo or naranjillo). We characterized three additional *S. quitoense* trichome expressed enzymes, and found that virus induced gene silencing of each caused changes in acylinositol accumulation. Surprisingly, the *in vitro* triacylinositol products of these enzymes are distinct from those that accumulate *in planta*. These enzymes, nonetheless, provide an opportunities to test the biological impact and properties of these triacylinositols *in vitro*.

## Introduction

Plant specialized metabolites are taxonomically-restricted small molecules that mediate a variety of interactions with the environment (Howe and Jander, 2008; Mithöfer and Boland, 2012; Massalha et al., 2017). These include protection from deleterious microbes and herbivores (Howe and Jander, 2008; Ahuja et al., 2012) and abiotic stresses such as UV-B (Landry et al., 1995), as well as interactions with beneficial bacterial and fungal partners, including symbionts (Massalha et al., 2017) and pollinators (Stevenson et al., 2017). Despite their diverse structures and sometimes complex biosynthesis, specialized metabolites are often made from simple components such as amino acids, sugars, and fatty acids as well as core metabolic intermediates.

Acylsugars are a class of specialized metabolites produced across the Solanaceae family (Moghe et al., 2017; Fan et al., 2019). These molecules contain a sugar core modified by esterification by one or more acyl groups. The acyl chains are derived from fatty acid and amino acid metabolism with an array of intraspecific diversity (Kim et al., 2012; Ning et al., 2015; Fan et al., 2020; Mandal et al., 2020; Landis et al., 2021); they range from two to twenty carbons in length with straight or branched configurations (Herrera-Salgado et al., 2005; Fan et al., 2019). Many acylsugars contain a sucrose core (King et al., 1986; Maldonado et al., 2006; Ghosh et al., 2014; Liu et al., 2017; Moghe et al., 2017), while others produce acylhexoses, including glucose and inositol (Burke et al., 1987; King and Calhoun, 1988; Matsuzaki et al., 1989). One example is the wild tomato *Solanum pennellii* (LA0716), which uses an invertase-like enzyme to convert acylsucroses to acylglucoses (Leong et al., 2019; Lybrand et al., 2020). Another is the black nightshade *Solanum nigrum*, an old-world Solanaceae species that uses a phylogenetically distinct invertase-like enzyme to synthesize acylglucoses (Lou et al., 2021).

Enzymes of various classes were demonstrated to play roles in acylsugar biosynthesis. The sugar acylating enzymes are clade III BAHD (named after the first four enzymes of this class) acyltransferases known as acylsugar acyltransferases (ASATs), which modify sugar cores using Coenzyme A thioesters as acyl group donors (Schilmiller et al., 2015; Fan et al., 2016; Moghe et al., 2017; Nadakuduti et al., 2017). Previous publications described characterization of ASATs across the Solanaceae: in cultivated (*Solanum lycopersicum*) and wild tomato species (i.e. *S. pennellii*) (Schilmiller et al., 2012; Schilmiller et al., 2015; Fan et al., 2017), *S. nigrum* (Lou et al., 2021)*, Petunia axillaris* (Nadakuduti et al., 2017), and *Salpiglossis sinuata* (Moghe et al., 2017). Trichome-expressed variants of primary metabolic enzymes, including isopropylmalate synthase, enoyl-CoA hydratase and CoA ligase provide ASAT acyl CoA substrates in cultivated tomato and *S. quitoense* (Ning et al., 2015; Fan et al., 2020). In addition, acylsugar hydrolases that remove acyl chains were identified, but their biosynthetic roles are as yet undefined (Ghosh et al., 2014; Schilmiller et al., 2016). These enzymes play distinct, but important, roles in acylsugar biosynthesis.

The presence of acylinositols in *S. quitoense*, *S. nigrum*, and *Solanum lanceolatum* offers an opportunity to study acylsugar core diversity in the *Solanum* clade (Herrera-Salgado et al., 2005; Leong et al., 2020). While members of the Leptostemonum clade – the largest monophyletic clade in the *Solanum* genus – were not thoroughly explored in a previous survey of acylsugars across the Solanaceae family (Moghe et al., 2017), *S. quitoense* and *S. lanceolatum* both produce acylated inositols. *S. quitoense* acylinositols with acyl chains of two, ten, and twelve carbons on a *myo-*inositol core are the major acylsugars (Figure S1), along with less abundant acylinositol glycosides (Hurney, 2018; Leong et al., 2020). *S. lanceolatum* acylinositol glycosides primarily have an acyl chain from twelve to twenty carbons (Herrera-Salgado et al., 2005). Only one acylsugar acyltransferase has been described in either of these species: a triacylinositol acetyltransferase (TAIAT) that synthesizes tetraacylinositols in *S. quitoense* (Leong et al., 2020). This system presents an opportunity to understand how acylsugar pathways have diverged in the *Solanum* clade.

We sought to identify other BAHD acyltransferases involved in acylinositol biosynthesis based upon characteristics of known ASATs. These include phylogenetic relationships, presence of conserved domains and enriched glandular trichome expression (Leong et al., 2020). Here we report the identification of three tomato ASAT homologs; SqASAT1H, SqASAT3H and a homolog of ASAT4 (SqASAT4H). Consistent with roles in acylinositol metabolism, virus-induced gene silencing (VIGS) of each candidate caused alteration of acylinositol quantity or acylation pattern. Screening recombinant ASATs *in vitro* demonstrated that ASAT1H acylated *myo*-inositol using nC10- and nC12-CoA, which are chains that match *in planta* products (n signifying straight chain). The ASAT1H substrate affinity for sugar and acyl-CoAs matches other ASATs. However, we found that ASAT1H *in vitro* acylation position does not match *in planta* products. This may be due to rearrangement of the monoacylinositol, which has been previously described for monoacylsucroses (Fan et al., 2016; Lou et al., 2021). ASAT4H and ASAT3H acylate the original monoinositol and subsequent diacylinositol to generate di- and triacylinositols, respectively. Contrary to expectation, the resulting triacylinositols do not co-elute with the *in planta* product.

## Results and Discussion

### Candidate gene identification

We sought acylinositol biosynthetic enzymes by screening for trichome-expressed BAHD acyltransferases that are phylogenetically related to characterized ASATs (Figure 1). We employed the same criteria used to identify TAIAT (Leong et al., 2020): seeking candidates with ≥500 reads in trichome RNAseq samples, predicted to encode proteins 400-500 amino acids in length with the canonical BAHD motifs HXXXD and DFGWG present in proper orientation (D’Auria, 2006). Three transcripts assemblies met these criteria: c39979_g2_i1, c38687_g1_i1, and c8981_g1_i1. All three candidates were expressed and enriched in *S. quitoense* trichomes relative to the trichome-denuded petiole tissue (Table 1) (Moghe et al., 2017). The first candidate—*c39979_g2_i1* (named ASAT1H below)—is a homolog of both the first enzyme of cultivated tomato acylsucrose biosynthesis, *Sl-ASAT1*, and the *Solanum lycopersicum* chromosome 7 *Sl-ASAT1* paralog *Solyc07g043670* (Moghe et al., 2017; Leong et al., 2020). The second candidate is the Sl-ASAT3 homolog *c8981_g1_i1* (referred to below as ASAT3H), while *c38687_g1_i1* (ASAT4H) is a paralog of the recently described triacylinositol acetyltransferase (TAIAT) (Leong et al., 2020).

**Figure 1.**
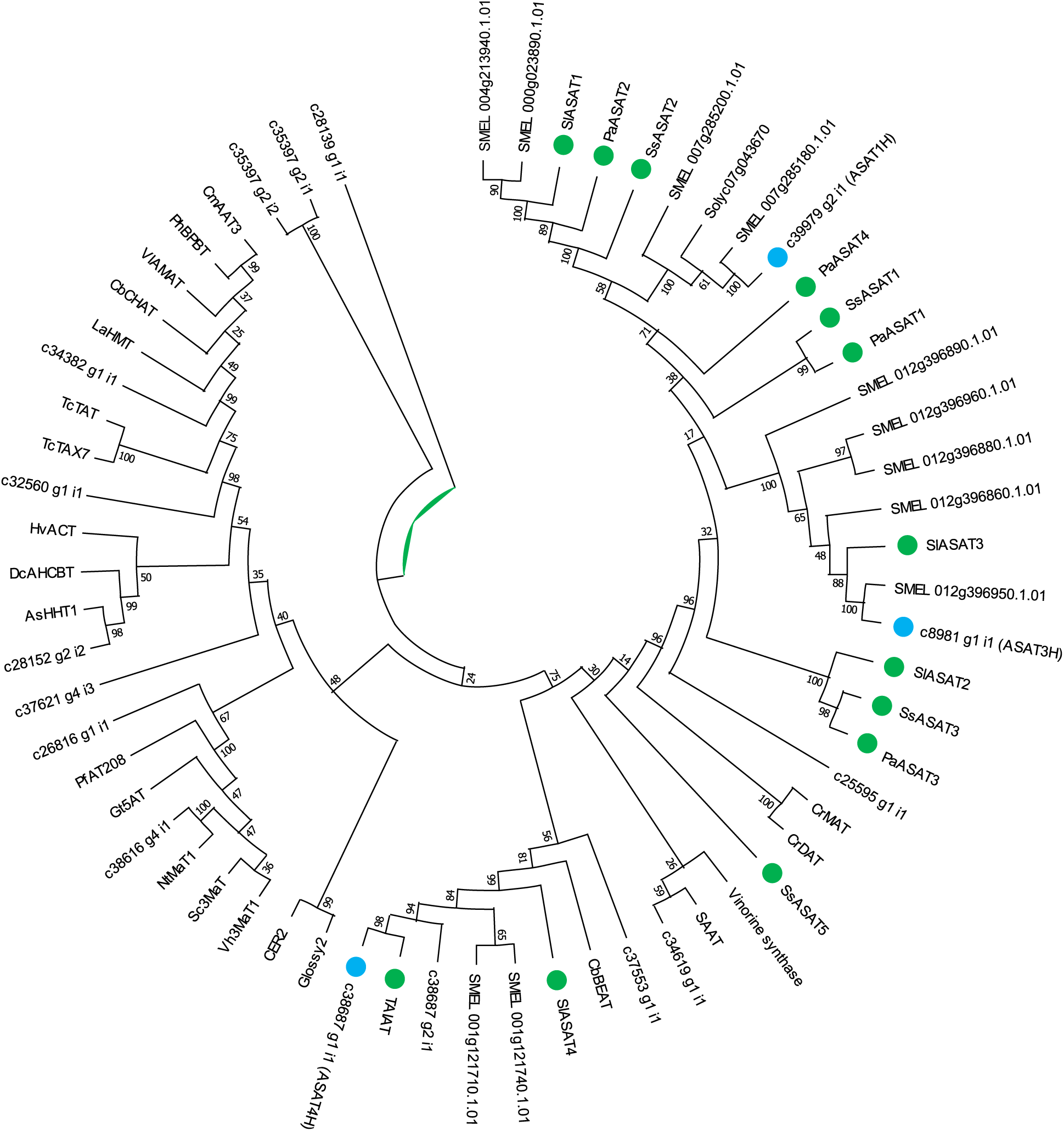
Phylogenetic analysis of known BAHD acyltransferases and *S. quitoense* ASAT candidates. Previously characterized ASATs are marked by green circles, while the enzymes described in this study are marked by blue circles. Protein sequences were aligned using MUSCLE with default parameters in MEGA X.

**Table 1.**
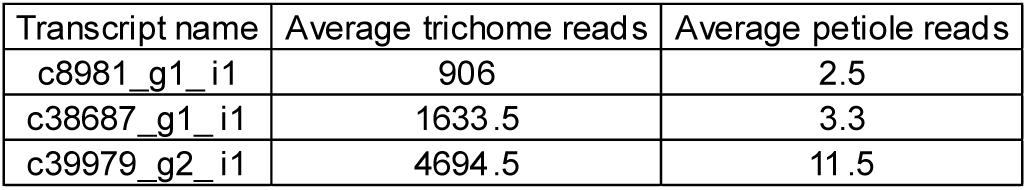
Trichome and shaved petiole RNA-seq data for tested *S. quitoense* ASAT candidates. Data are derived from Moghe et al. (2017).

### *In vivo* analysis of acylsugar acyltransferase candidates

A previously-developed *S. quitoense* VIGS method was employed to test the impact of downregulating the expression of these candidates *in vivo* (Figure S2). Qualitatively similar results were obtained for each of the two distinct fragments targeting ASAT1H, ASAT3H or ASAT4H transcripts.

VIGS of all three candidates yielded acylsugar phenotype perturbations. Targeting ASAT1H, ASAT3H, and ASAT4H individually caused reductions in the four major acylinositols (Figure 2A and B, Table S1 and Table S2). This phenotype is reminiscent of results from CRISPR knockout lines of ASATs in *S. lycopersicum and S. pennellii*, as well as VIGS of *Petunia axillaris*, *Salpiglossis sinuata*, *S. nigrum* (Lou et al., 2021) and *S. quitoense* ASATs (Schilmiller et al., 2015; Fan et al., 2016; Moghe et al., 2017; Nadakuduti et al., 2017; Leong, 2019). qRT-PCR analysis confirmed reductions in candidate transcript abundance in target plants relative to the controls, with three of six constructs significant at *p* < 0.05 (Figure S3 and S4). The decreased acylsugar accumulation suggests a role in acylinositol biosynthesis and the case for ASAT1H and ASAT3H is bolstered by a lack of other significant paralogs or homology to other candidates that could result in cross-silencing. Targeting ASAT4H—but not ASAT1H or ASAT3H—yielded an increased ratio of monoacetylated tri- to diacetylated tetra-acylinositols (Figure S5, Table S1, and Table S2). We hypothesized that the similarity between TAIAT and ASAT4H (93% nucleotide identity) could cause cross-silencing (Table S3, Figure S6); this was validated by qRT-PCR analysis, which revealed that targeting ASAT4H caused reduced TAIAT transcript abundance (Figure S4). Thus, TAIAT cross-silencing could have contributed to the reduction in tetraacylinositols that contained two acetate esters. The perturbation of acylinositol accumulation led us to further characterize these candidates using *in vitro* biochemistry.

**Figure 2.**
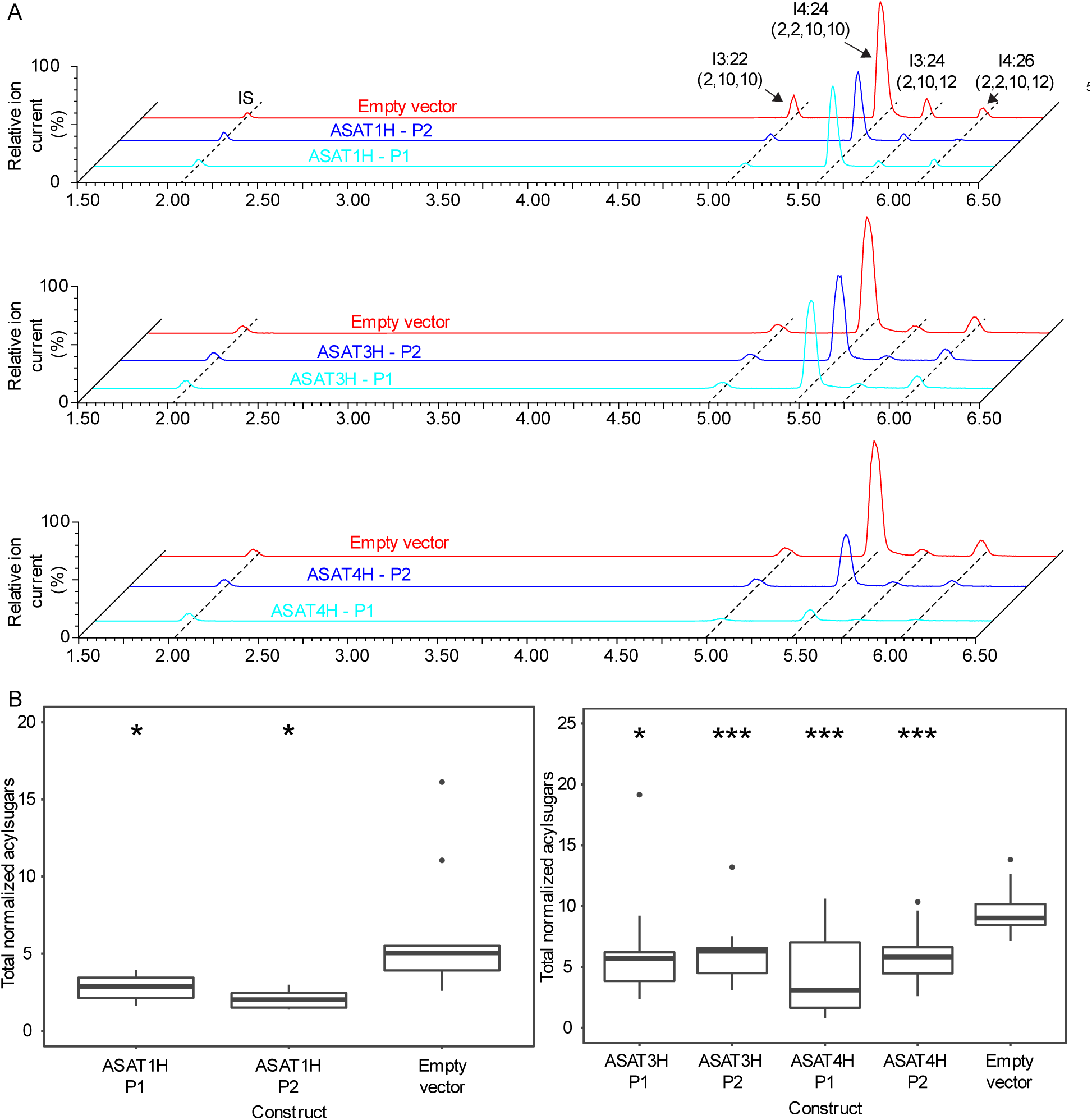
VIGS analysis of acylinositol biosynthesis candidates. A, LC-MS analysis of acylsugars in ASAT1H, ASAT3H, and ASAT4H-targeted plants from similar sized leaves (< 0.6 mg difference). The combined extracted ion chromatogram shows minute 1.50 to 6.50 of a 7 minute method with telmisartan (internal standard), I3:22 (2,10,10), I4:24 (2,2,10,10), I3:24 (2,10,12), and I4:26 (2,2,10,12). Chromatograms combined signals: [M-H]^-^ for telmisartan and [M+formate]^-^ for the acylsugars. The *m/z* values of compounds included in this figure: telmisartan (internal standard), *m/z* 513.23; I3:22 (2,10,10), *m/z* 577.35; I4:24 (2,2,10,10), *m/z* 619.37; I3:24 (2,10,12), *m/z* 605.4; I4:26 (2,2,10,12), *m/z* 647.4, each with a mass window of 0.05 Da. B, Total acylsugar boxplots in ASAT1H, ASAT3H, and ASAT4H VIGS samples. Welch’s two-sample t-test was used for pairwise statistical analysis to empty vector controls. Whiskers represent minimum and maximum values less than 1.5 times the interquartile range from the first and third quartiles, respectively. Values outside of this range are depicted using black dots. *P < 0.05, **P < 0.01, and ***P < 0.001. For ASAT1H, ASAT3H, ASAT4H-targeted plants (n = 14-15), for ASAT1H empty vector plants (n = 9), for ASAT3H and ASAT4H empty vector plants (n = 14).

### *In vitro* analysis of ASAT1H

ASAT1H emerged as the top candidate for the first step in acylinositol biosynthesis based on its amino acid similarity (44% amino acid identity) and phylogenetic relationship to Sl-ASAT1—the enzyme that acylates sucrose in cultivated tomato acylsucrose biosynthesis (Figure 1, Table S3) (Fan et al., 2016)— as well as VIGS-mediated reduction in total acylinositols. We used *in vitro* biochemistry to determine whether an *E. coli*-expressed ASAT1H acylates *myo*-inositol using nC10 and nC12-CoA as substrates. These acyl donors were tested because *S. quitoense* trichome acylsugars contain acylinositols esterified with nC10 and nC12 acyl groups (Leong et al., 2020). To minimize rearrangement of the primary products, assays were performed at pH 6 (Fan et al., 2016). Indeed, ASAT1H catalyzed formation of monoacylated inositol using both nC10 and nC12-CoAs and inositol as substrates (Figure 3A). Mass spectrometric analysis using negative-ion mode collision-induced dissociation yielded *m/z* 171.14 or 199.17 fragment ions, values consistent with negatively charged carboxylates of nC10 or nC12 acyl chains, respectively (Figure S7).

**Figure 3.**
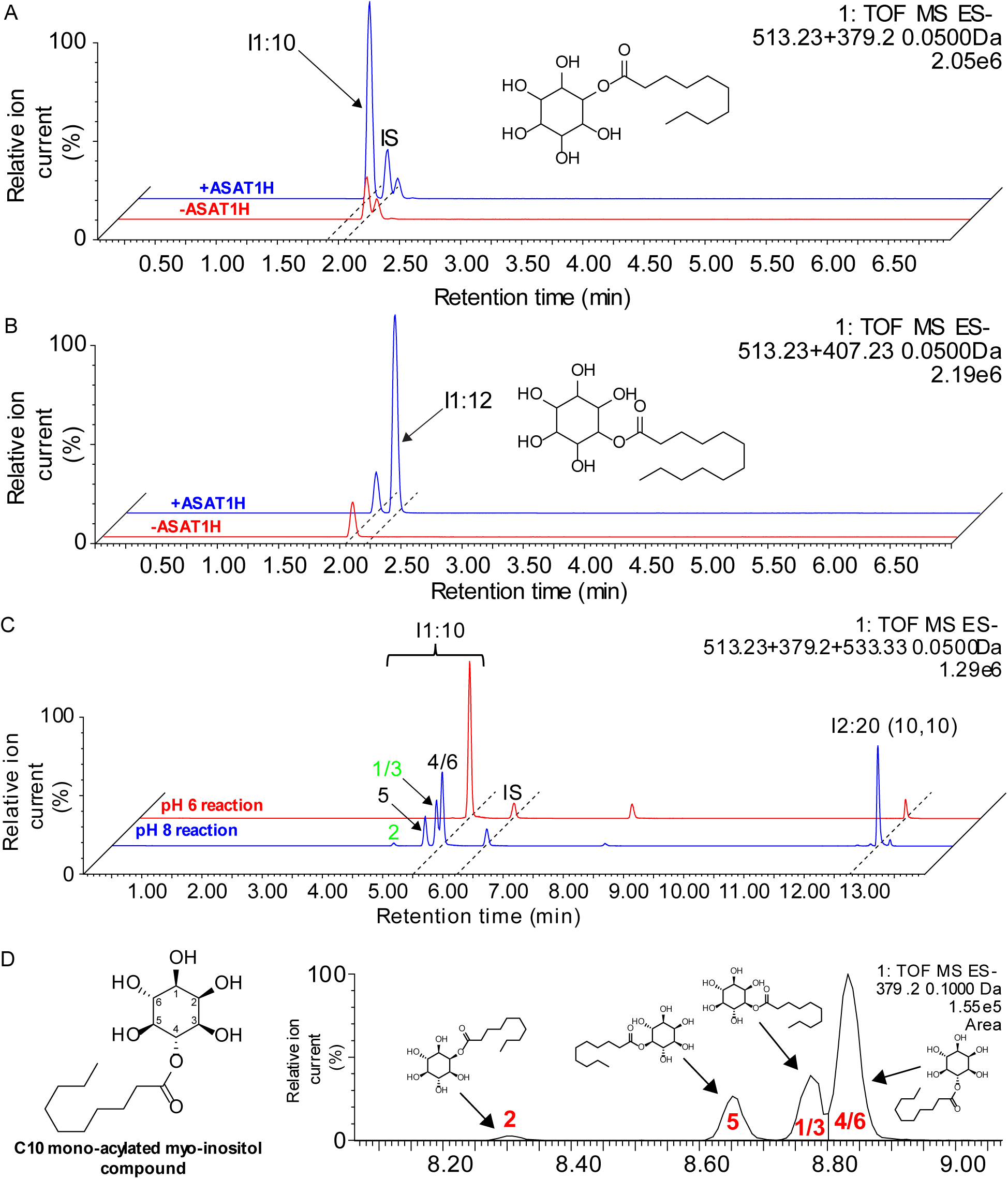
ASAT1H *in vitro* assay product characterization. LC-MS analysis of ASAT1H *in vitro* assay using *myo*-inositol and A, nC10-CoA or B, nC12-CoA. The combined extracted ion chromatogram includes telmisartan (internal standard), I1:10, or I1:12. C, LC-MS analysis of ASAT1H assays at pH 6 (red) and 8 (blue). The combined extracted ion chromatogram includes telmisartan (internal standard), I1:10, and I2:20 (10,10). I1:10 isomers are labeled and *in vivo* positions of medium acyl chains are green. D, NMR-derived structure of ASAT1H I1:10 product (left). The extracted ion chromatogram shows relative elution order of monoacylinositol isomers using reverse phase chromatography on 14 minute method with a 0.1 Da mass window (right). The *m/z* values of the compounds in this figure: telmisartan (internal standard), *m/z* 513.23; I1:10, *m/z* 379.20; I1:12, *m/z* 407.23; I2:20 (10,10), *m/z* 533.33, each with a mass window of 0.05 Da unless otherwise described. All acylsugars are shown as formate adducts.

We posited that if ASAT1H is involved in acylinositol biosynthesis, it would have donor and acceptor substrate affinities similar to other ASATs (Schilmiller et al., 2015; Fan et al., 2016). Indeed, the apparent *K*_m_ for *myo*-inositol with nC10-CoA was 4.4 ± 1.1 mM (95% confidence interval using standard error) (Figure S8 and Table S4), which is similar to the Sl-ASAT1 apparent *K*_m_ for sucrose of 2.3 mM (Fan et al., 2016). The apparent *K*_m_ of ASAT1H for nC10-CoA and nC12-CoA with *myo*-inositol were 9.5 ± 2.4 µM and 16.6 ± 6.6 µM, respectively. These match the previously reported values for acyl-CoAs of 2-50 µM for Sl-ASAT1, SlASAT2, and ASAT3 enzymes from multiple species (Schilmiller et al., 2015; Fan et al., 2016). Taken together, the ASAT1H apparent *K*_m_ donor and acceptor substrate results match other ASATs, consistent with a possible role in acylinositol biosynthesis.

We resolved the I1:10 monoacylinositol product of reaction of nC10-CoA and *myo*-inositol using NMR and compared the acylation position to *in planta* acylsugars. I1:10 describes a *myo*-inositol core with one acyl chain of ten carbons; with more than one acyl chain, the numbers of carbon atoms of individual chains are included in parentheses (for example, I3:22 (2,10,10). ASAT1H generated a single I1:10 product in reactions run at pH 6, a condition that previously yielded monoacylsucrose esterified at the 4-position with tomato SlASAT1 (Figure 3B) (Fan et al., 2016). Analysis of the purified I1:10 NMR spectra revealed the product to be acylated at the *myo*-inositol 4/6 position (Figure 3C and Table S5). This result was unexpected because NMR-resolved structures of *S. quitoense* acylsugars contain nC10 or nC12 acyl chains at the 2- and 1-/3- positions of *myo*-inositol, without evidence of 4-position acylation (Hurney, 2018; Leong et al., 2020). Note acylation at that *myo-*inositol positions 1/3 and 4/6 are mirror plane symmetrical, and thus indistinguishable by NMR and reverse-phase LC. These results show that the *in vitro* I1:10 does not match *in planta* acylsugars.

In contrast to the pH 6 assay product, we found a mixture of four chromatographically distinct I1:10 *myo*-inositol isomers in reactions at pH ≥7 (Figure 3C). ^1^H-NMR analysis revealed four monoacylinositol positional isomers at relative peak integration of 2:15:19:64, correlating well with the LC-MS peak area ratios of approximately 2:15:22:61 (Figure 3C, Figure S9.A and B, and Table S6). Therefore, we inferred that the monoacylinositol I1:10 esterified at position 2 eluted first, followed by that at position 5, position 1/3 and finally position 4/6.

As ASAT1 *in vitro* enzyme assays run at pH 6 and pH 8 produced different product profiles (Figure 3C), we tested the hypothesis that rearrangement of the 4-position monoacylinositol primary product to position 2 or 1/3 contributes to *in vivo* biosynthesis. If rearrangement is occurring, the isomers esterified at positions other than 4 should increase with reaction time at pH 8. Indeed, the other isomers, absent at pH 6, increased in abundance over the course of an hour after a shift to pH 8, along with a peak matching the *m/z* of the diacylated compound I2:20 (10,10) (*m/z* 533.33) (Figure S10). Some of these isomers matched the position of medium chains from *in planta* acylsugars. Rearrangement did not require presence of the enzyme, as incubation of purified 4-position ester I1:10 product without enzyme at pH 8 for 60 minutes at 30°C yielded rearranged I1:10 isomers (Figure S11). We asked whether the rearrangement of the R4 acylchain was inter- or intramolecular by comparing rate and extent of rearrangement following serial dilution. First-order kinetics were obtained, consistent with intramolecular rearrangement: up to a nine-fold dilution did not impact product formation over 30-60 minutes (Figure S12). Taken together, these results indicate that a single R4 monoacylinositol is formed *in vitro* at pH 6, and rearranges non-enzymatically at pH 8. This is reminiscent of the Sl-ASAT1 and SnASAT1 R4 monoacylated sucrose product rearrangement to form R6 monoacylated sucrose at ≥ pH 7 (Fan et al., 2016) (Lou et al., 2021). While the absence of acylsugars with acyl groups in the 6-position in tomato and *S. nigrum* strongly suggests that rearrangement is not part of acylsugar biosynthesis in these plants (Fan et al., 2016; Lou et al., 2021), we cannot rule out a role of intramolecular rearrangement in acylinositol metabolism.

### ASAT3H and ASAT4H produce triacylinositols that differ from the *in vivo* products

We performed *in vitro* assays to ask whether ASAT3H or ASAT4H have activities consistent with roles in acylinositol biosynthesis. We used a combined reaction containing *E. coli*-expressed His-tagged ASAT1H, ASAT4H, and ASAT3H with C2-CoA, nC10-CoA and *myo*-inositol substrates. These acyl-CoA donors were used because those acyl chains are found in *S. quitoense* acylinositols. This assay produced an I3:22 (2,10,10) peak at pH 6 or 8 (Figure 4A, Figure S13). MS fragmentation of the I3:22 (2,10,10) *in vitro* product matched the major *in planta* acylinositol (Figure S14), but the two compounds did not co-elute; this is consistent with acylation positions different from those of the *in vivo* metabolite (Figure 4B). The difference in acylation positions indicates that *S. quitoense* acylinositol biosynthesis requires more than consecutive acylations of *myo*-inositol by the ASAT1H, ASAT3H, and ASAT4H enzymes.

**Figure 4.**
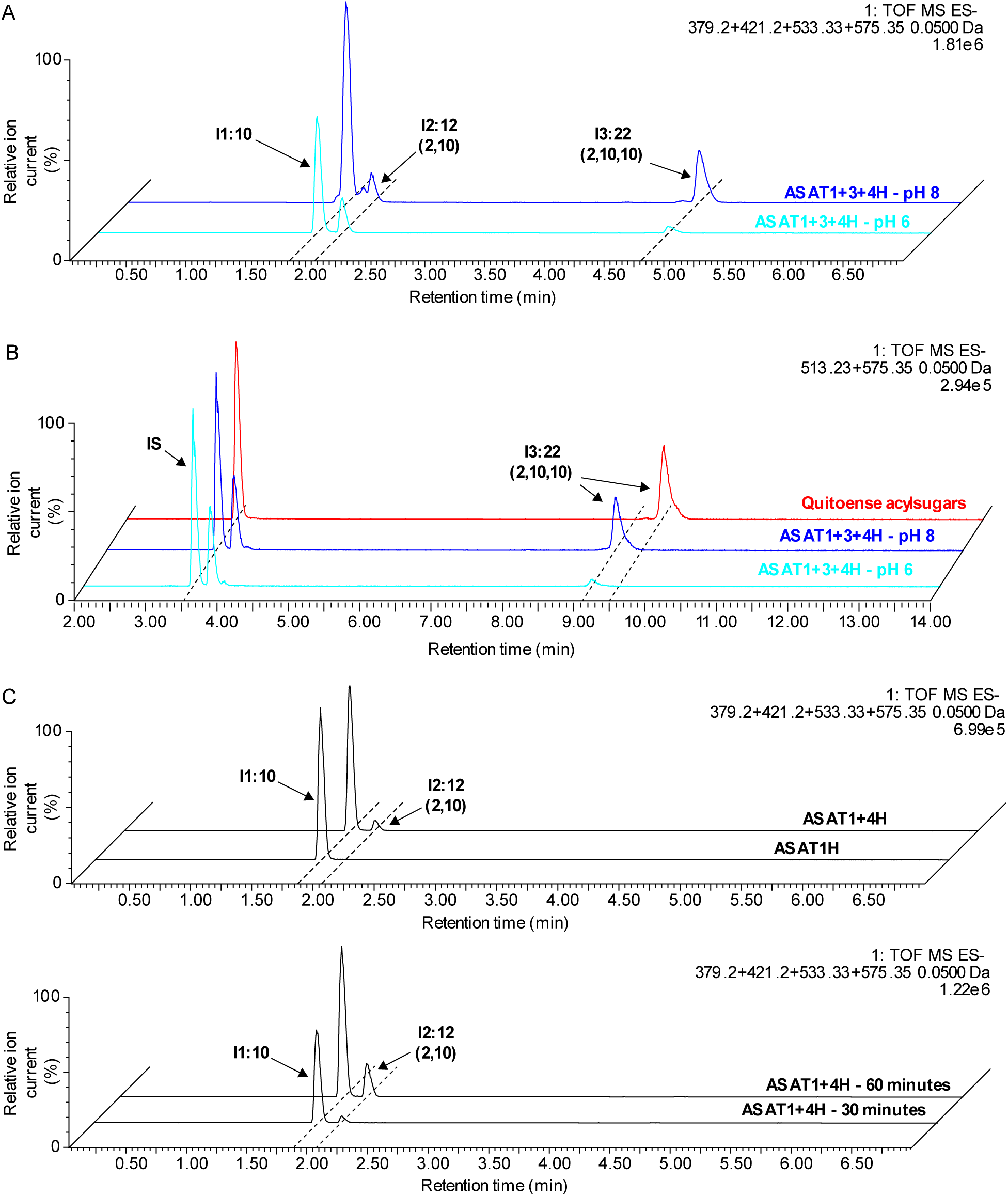
ASAT1H, ASAT3H, and ASAT4H *in vitro* assay product characterization. A, LC-MS analysis of combined assays with ASAT1H, ASAT3H, and ASAT4H using *myo*-inositol, nC10-CoA and C2-CoA at pH 6 (light blue) or 8 (blue). The combined extracted ion chromatogram includes I1:10, I2:12 (2,10), I2:20 (10,10), and I3:22 (2,10,10). B, LC-MS analysis of *S. quitoense* acylsugars (red), and ASAT1H, ASAT3H, and ASAT4H combined assays using *myo*-inositol, nC10-CoA and C2-CoA at pH 6 (light blue) or pH 8 (blue) using 21 minute method. The combined extracted ion chromatogram includes telmisartan (internal standard) and I3:22 (2,10,10). C, LC-MS analysis of ASAT1H and ASAT4H combined assays using *myo*-inositol, nC10-CoA and C2-CoA at pH 6. Combined extracted ion chromatogram of a 30 minute reaction includes I1:10, I2:12 (2,10), I2:20 (10,10), and I3:22 (2,10,10) (top) with reaction progress at 30 and 60 minutes (bottom). The *m/z* values of the compounds included in this figure: telmisartan (internal standard), *m/z* 513.23; I1:10, *m/z* 379.20; I1:12, *m/z* 407.23; I2:20 (10,10), *m/z* 533.33; I3:22 (2,10,10), *m/z* 575.35, all with a mass window of 0.05 Da. All acylsugars are shown as formate adducts.

The one pot combined reaction did not determine the relative order of action of ASAT4H and ASAT3H. To test the hypothesis that ASAT4H acetylates the I1:10 ASAT1H *in vitro* product, we combined ASAT1H and ASAT4H in a reaction with *myo*-inositol, C2-CoA and nC10-CoA at pH 6, yielding an I2:12 (2,10) product (Figure 4C, Figure S15). Acetylation of I1:10 by ASAT4H was shown to occur with sequential assays containing C2-CoA, nC10-CoA and *myo*-inositol with ASAT1H followed by heat inactivation and addition of ASAT4H, C2-CoA and nC10-CoA yielded the I2:12 (2,10) product (Figure S16). In contrast, the reciprocal order – ASAT4H followed by ASAT1H – failed to yield I2:12 (2,10). These results indicate that ASAT1H acylates *myo*-inositol with nC10-CoA at position R6, followed by acetylation by ASAT4H.

The cross-silencing of the other SlASAT4 homolog, SqTAIAT, by VIGS directed against ASAT4H (Figure S4) led us to ask whether it would also use I1:10^R6^ as acyl acceptor. However, in contrast to ASAT4H, the combination of ASAT1H and TAIAT did not yield I2:12 (2,10) (Figure S17). While the SqASAT4H acetyltransferase activity matches the other ASAT4 clade proteins Sl-ASAT4, SsASAT5, and TAIAT (Schilmiller et al., 2012; Leong et al., 2020), these other enzymes act on acceptor substrates that are tri- or tetra-acylated. ASAT4H is thus far unique in acetylating a monoacylated product *in vitro* (Fan et al., 2016; Moghe et al., 2017; Leong et al., 2020).

The results with a one pot assay containing three enzymes described above suggest that ASAT3H adds an nC10 acyl chain to I2:12 (2,10) to produce I3:22 (2,10,10). We used sequential assays to test this: two enzymes were added together with C2-CoA, nC10-CoA and *myo*-inositol, followed by heat inactivation, and the addition of the third enzyme with more C2-CoA and nC10-CoA. The reaction with ASAT1H and ASAT4H followed by ASAT3H yielded both I2:12 (2,10) and I3:22 (2,10,10) consistent with ASAT3H catalyzing a third acylation (Figure S18). The reciprocal reaction with ASAT1H and ASAT3H followed by ASAT4H produced I2:12 (2,10) (presumably from consecutive reaction of ASAT1H and ASAT4H), but not I3:22 (2,10,10) (Figure S18). Taken together, these results indicate that ASAT3H can add an nC10 acyl chain to I2:12 (2,10) to make I3:22 (2,10,10), but does not acylate I1:10 to yield I2:20 (10,10). This is analogous to tomato acylsugar biosynthesis, where Sl-ASAT3 acylates diacylsucroses to yield triacylsucroses (Schilmiller et al., 2015; Fan et al., 2016).

## Conclusions

*S. quitoense* acylinositol biosynthesis is interesting due to the sugar core difference from previously characterized acylsugar pathways (Fan et al., 2016; Fan et al., 2017; Moghe et al., 2017; Nadakuduti et al., 2017). We hypothesized that acylinositol biosynthesis is evolutionarily related to acylsucrose biosynthesis in other Solanaceae species (Fan et al., 2016; Moghe et al., 2017; Leong et al., 2020). An *S. quitoense* triacylinositol acetyltransferase (TAIAT) was previously shown to acetylate triacylinositols yielding tetracylinositols (Leong et al., 2020). In this study we describe characterization of three candidates (ASAT1H, ASAT4H, and ASAT3H) that are homologous to previously described ASATs (Figure 1)(Schilmiller et al., 2012; Schilmiller et al., 2015; Fan et al., 2016; Moghe et al., 2017). These candidates are expressed and enriched in trichomes relative to shaved petiole tissue (Table 1) (Moghe et al., 2017). We used VIGS to test if these candidates are involved in *S. quitoense* acylinositol biosynthesis. VIGS results with both ASAT1H- and ASAT3H-targeted plants show reduced total acylsugars consistent with an early step in acylinositol biosynthesis (Fan et al., 2016; Moghe et al., 2017; Nadakuduti et al., 2017). Silencing ASAT4H yields reduced total acylsugars and increased ratio of tri- to tetraacylinositols. Interpreting the ASAT4H phenotype requires caution due to cross-silencing of TAIAT in the VIGS plants.

Results of *in vitro* biochemistry with these three candidates demonstrated ASAT-like activities with acceptor substrate specificities, but revealed products that are positional isomers of those detected from trichome extracts. ASAT1H catalyzes acylation of *myo*-inositol using nC10-CoA and nC12-CoA. It also possesses apparent *K*_m_ values similar to previously characterized ASATs. NMR analysis revealed that the acyl chain of the I1:10 product is not at a position on *myo*-inositol that matches the *in planta* acylsugars. *In vitro* assays revealed that the I1:10 product can rearrange at pH 8, even in the absence of an enzyme. Assays with ASAT1H, ASAT4H, and ASAT3H showed an I3:22 (2,10,10) product that did not co-elute with the *in planta* I3:22 (2,10,10). Further *in vitro* assays indicated that ASAT1H acylated *myo*-inositol with an nC10-CoA first, followed by ASAT4H-catalyzed acetylation and then nC10 acyl chain addition by ASAT3H.

We propose several possibilities that could reconcile the conflict between the *in vitro* and *in planta* results. First, our *in vitro* experiments could be missing one or more enzymes involved in acylation at the positions observed *in planta*, for example an acylsugar acyltransferase. This hypothesis could still include rearrangement at pH > 7. Second, another class of enzyme – such as a serine carboxypeptidase-like enzyme (SCPL) – could play a role in acylinositol biosynthesis. SCPLs play roles in transferring acyl chains in other biosynthetic pathways (Shirley et al., 2001; Stehle et al., 2008; Mugford et al., 2009). Acylsugar hydrolases (*S. quitoense* transcripts c33090_g1_i1/2 and c31412_g4_i1) are a third class of enzyme that could be involved *in vivo*: these remove acyl chains from acylsucroses and are trichome-enriched in *S. quitoense* (Schilmiller et al., 2016; Moghe et al., 2017). Perhaps edited acylsugars serve as substrates for other enzymes to generate the observed *in planta* product. Nonetheless, perturbation of the acylinositol phenotype due to VIGS of the BAHD enzymes strongly suggests that they are involved in acylinositol biosynthesis.

A third hypothesis is that *myo*-inositol is not the starting substrate for acylinositol biosynthesis. An analogous situation is seen in two *Solanum* species that make acylglucoses by acylating the disaccharide sucrose, followed by hydrolysis to acylglucose and fructose mediated by specialized invertases (Leong et al., 2019; Lou et al., 2021). In fact, *S. quitoense* accumulates acylated *myo*-inositol glycosides with sugar cores that could be the precursor of the *in planta* acylinositols (Hurney, 2018). *S. lanceolatum* is another species in the *Solanum* Leptostemonum clade that accumulates acylated inositol-glucose and inositol-xylose disaccharides (Herrera-Salgado et al., 2005). In the future, acylinositol accumulation in other *Solanum* species can be leveraged to understand how acylinositols are synthesized *in planta*.

While it is disappointing that the *in vitro* products of ASAT1H, ASAT3H, and ASAT4H do not match the esterification positions of *S. quitoense* acylinositols characterized from trichome extracts, they can be employed for further studies. For example, the structurally-distinct *in vitro*-produced acylinositols are useful for testing the efficacy of different acylsugars on insects and pathogens. Protein structure and function analysis could reveal details about BAHD catalysis. In addition, the BAHD acyltransferases could be used to engineer acylinositol biosynthesis into other plants or microorganisms to search for biological activities.

## Author contributions

All authors designed the original research; B.J.L., S.M.H., T.A., and P.D.F. performed the experiments; B.J.L., S.M.H., and R.L.L. wrote the manuscript; all authors edited the article.

## Material availability

Datasets and materials used in this study available upon reasonable request.

## Figure legends

**Figure S1.**
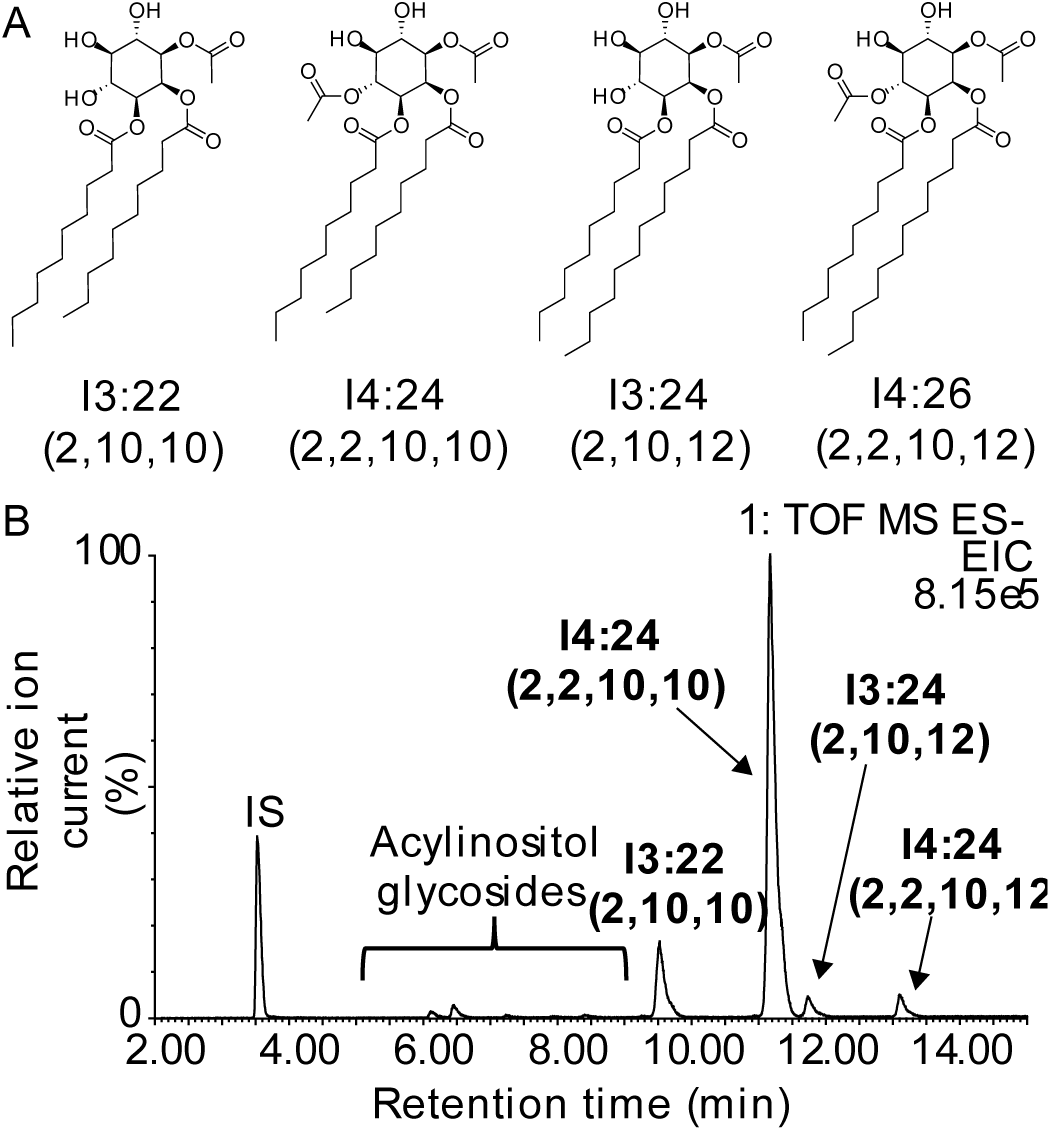
*Solanum quitoense* acylinositols. A, NMR-derived structures of previously identified *S. quitoense* tri- and tetraacylinositol monosaccharides. Note acylation at that *myo-*inositol positions 1/3 and 4/6 are mirror plane symmetrical, indistinguishable by NMR and reverse-phase LC-MS. B, LC-MS of previously characterized *S. quitoense* acylinositols (Leong et al., 2020) and acylinositol glycosides (Hurney, 2018). All acylsugars are shown as formate adducts.

**Figure S2.**
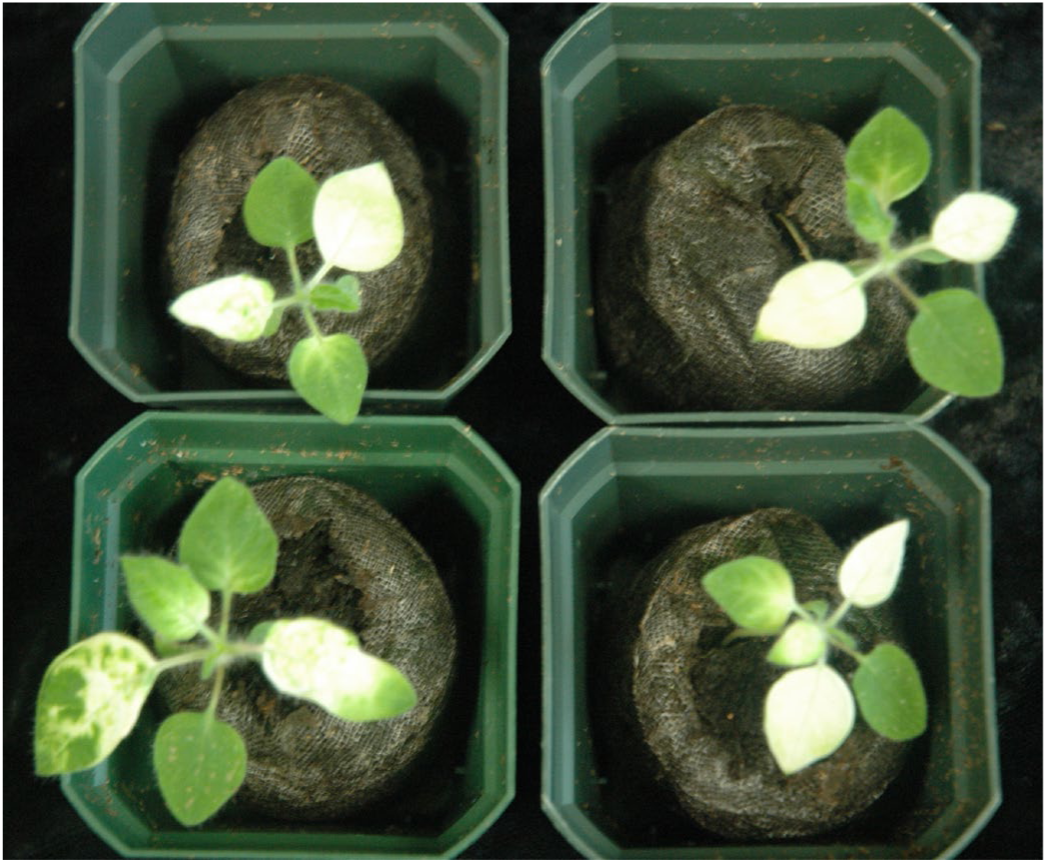
Whole plant phenotype of PDS-targeted *S. quitoense* plants.

**Figure S3.**
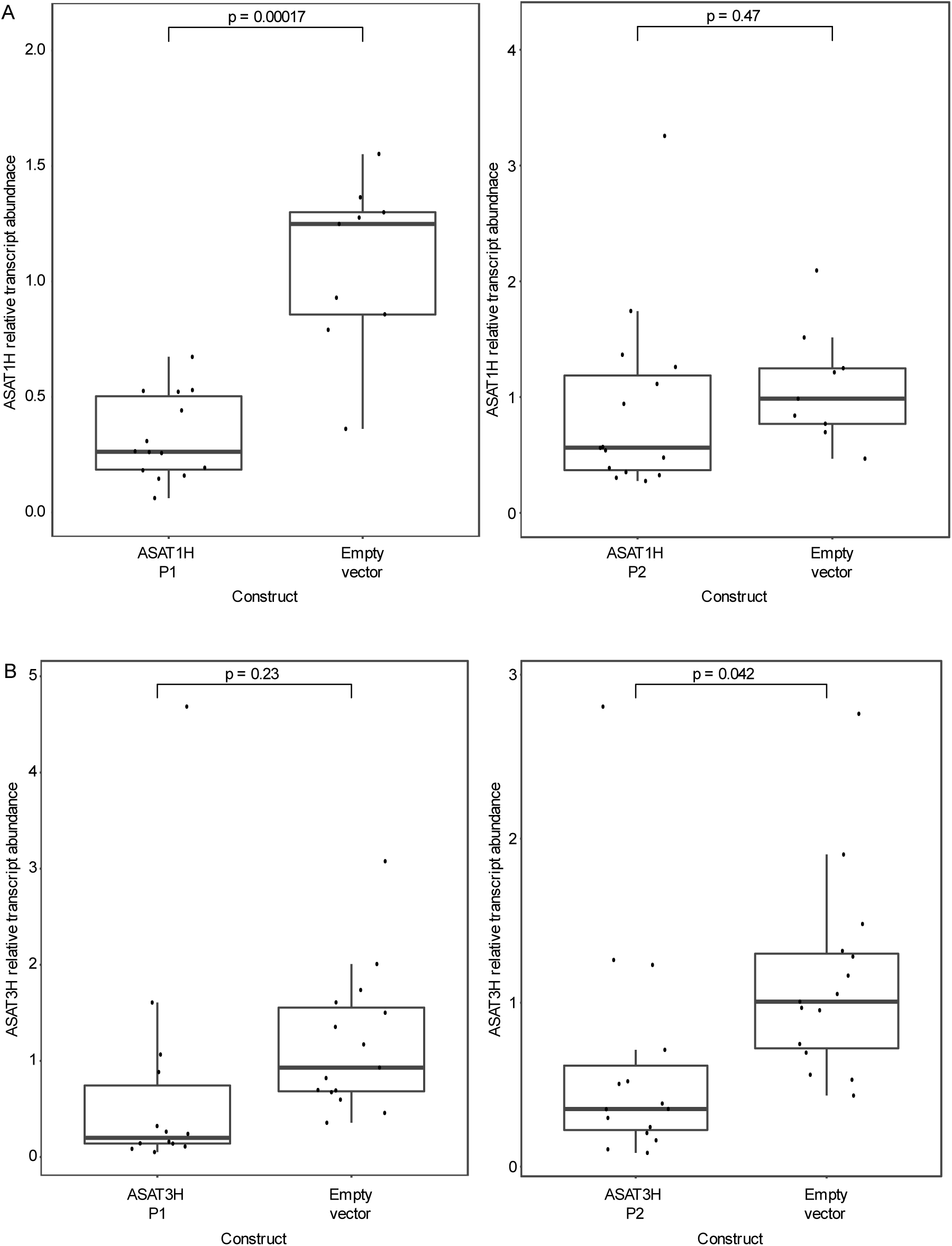
ASAT1H and ASAT3H relative transcript abundance in VIGS plants. A, ASAT1H relative transcript abundance plot in plants targeting different parts of the ASAT1H transcript using ASAT1H P1 construct (left) or ASAT1H P2 construct (right). B, ASAT3H relative transcript abundance plot in plants targeting different parts of the ASAT3H transcript using ASAT3H P1 construct (left), or ASAT3H P2 construct (right). Welch’s two-sample t-test was used for pairwise statistical analysis to empty vector controls. *P < 0.05, **P < 0.01, and ***P < 0.001. Whiskers represent minimum and maximum values less than 1.5 times the interquartile range from the first and third quartiles, respectively. For ASAT1H and ASAT3H-targeted plants, n = 15; for empty vector controls for ASAT1H and ASAT3H experiments n = 9 and 15, respectively with 3-4 technical replicates per biological sample.

**Figure S4.**
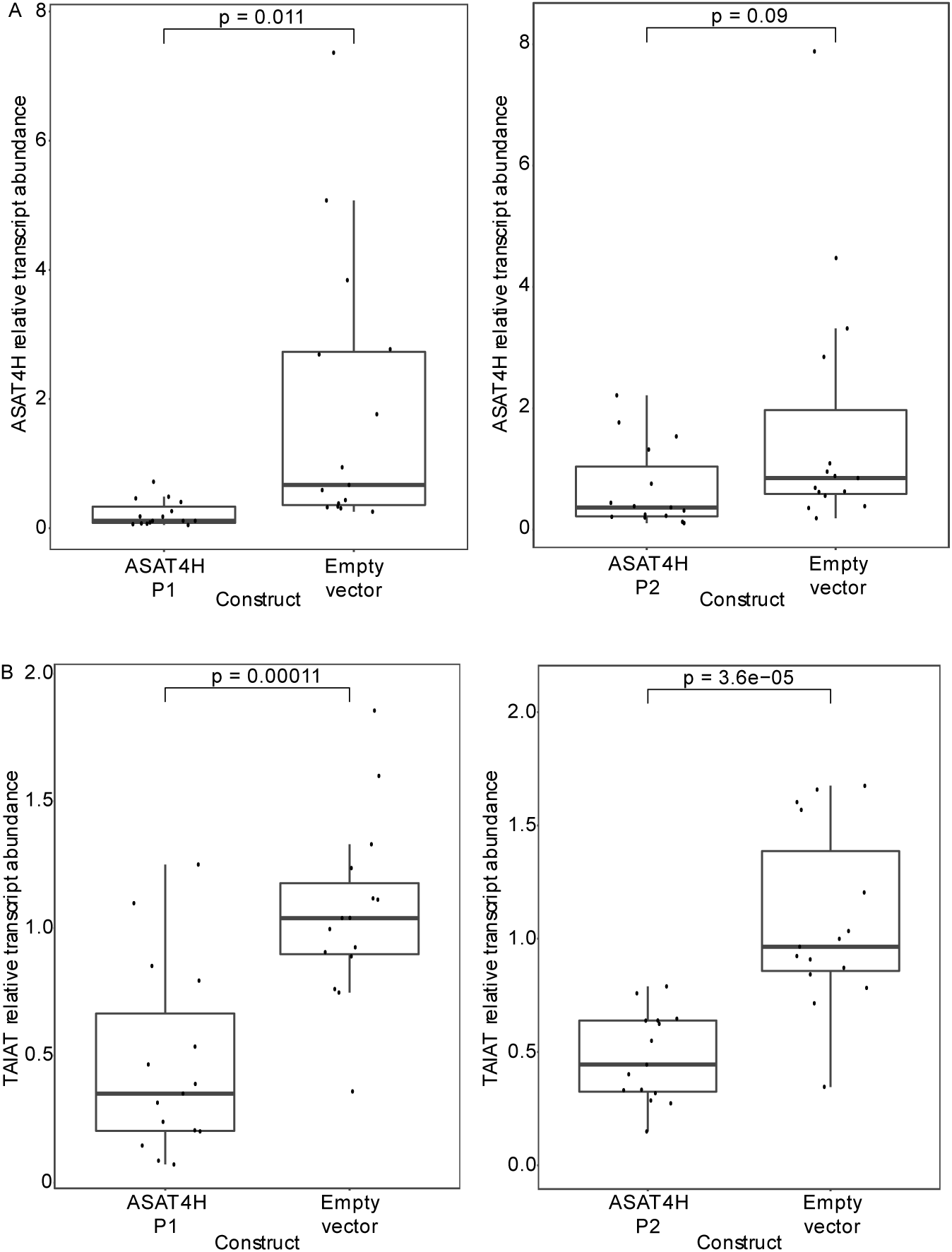
ASAT4H and TAIAT relative transcript abundance in ASAT4H-targeted VIGS plants. A, ASAT4H or B, TAIAT relative transcript abundance plot in plants targeting different parts of the ASAT4H transcript using ASAT4H P1 construct (left) or ASAT4H P2 construct (right). Welch’s two-sample t-test was used for pairwise statistical analysis to empty vector controls. *P < 0.05, **P < 0.01, and ***P < 0.001. Whiskers represent minimum and maximum values less than 1.5 times the interquartile range from the first and third quartiles, respectively. For ASAT4H-targeted plants and empty vector controls (n = 15) and 3-4 technical replicates per sample.

**Figure S5.**
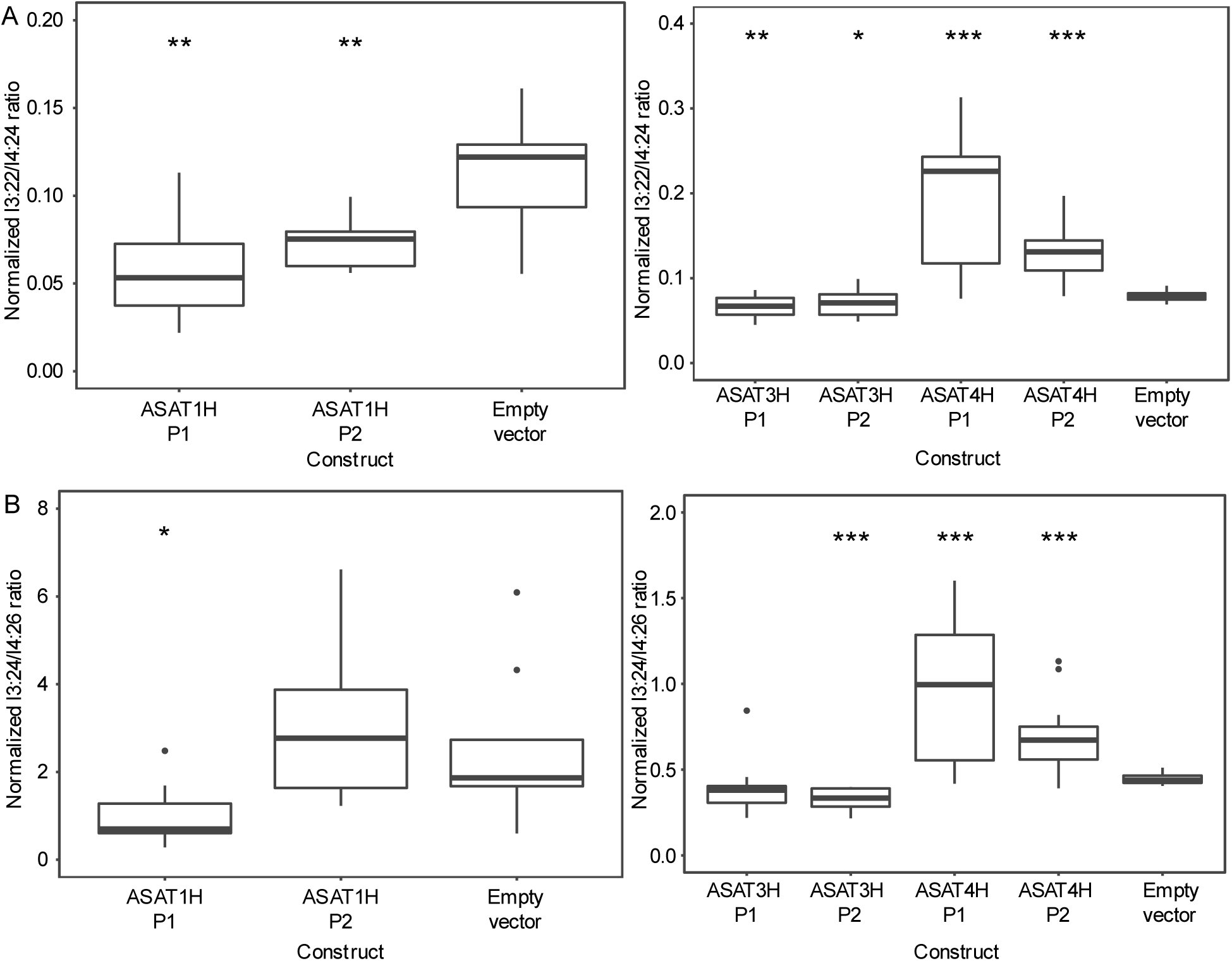
Analysis of tri- to tetraacylinositol ratios in VIGS plants. A, Ratio of I3:22 (2,10,10) to I4:24 (2,2,10,10) in ASAT1H, ASAT3H, and ASAT4H-targeted plants compared to empty vector control plants. B, Ratio of I3:24 (2,10,12) and I4:26 (2,2,10,12) in ASAT1H, ASAT3H and ASAT4H-targeted plants compared to empty vector controls. Values are the ratios of normalized acylinositol response. Normalized response represents peak areas (m+2 natural isotopologs of formate adducts) normalized to the internal standard (telmisartan) and dry weight of the extracted leaf. Welch’s two-sample t-test was performed between the test samples and the empty vector control plants to determine the p-value. *P < 0.05, **P < 0.01, and ***P < 0.001. Whiskers represent minimum and maximum values less than 1.5 times the interquartile range from the first and third quartiles, respectively. Values outside this range are depicted using black dots. For ASAT1H, ASAT3H, and ASAT4H-targeted plants (n = 13-15 per construct), for ASAT1H empty vector control plants (n = 9), for ASAT3H and ASAT4H empty vector control plants (n = 15). ASAT1H data are from a separate experiment from ASAT3H and ASAT4H data.

**Figure S6.**
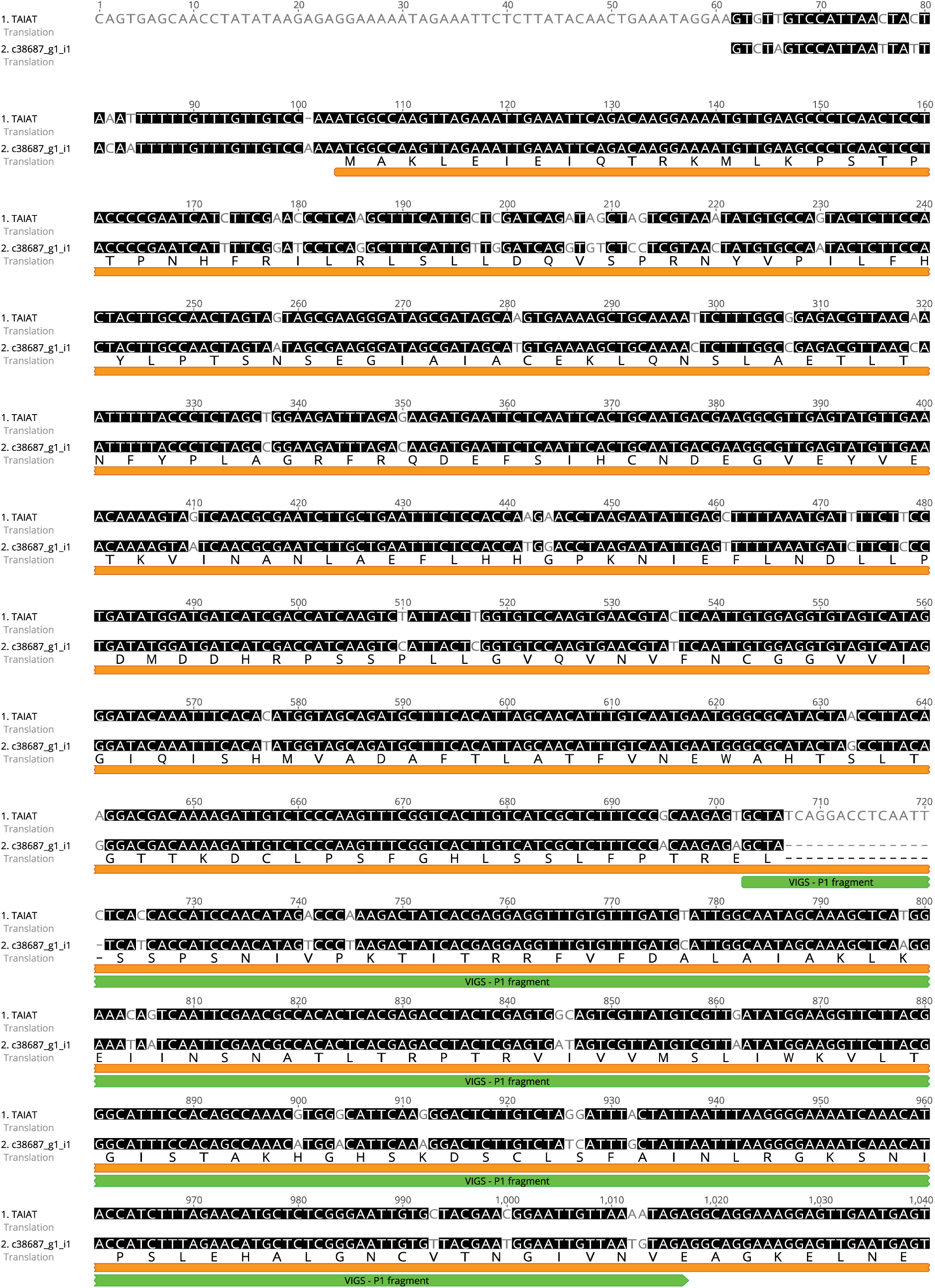
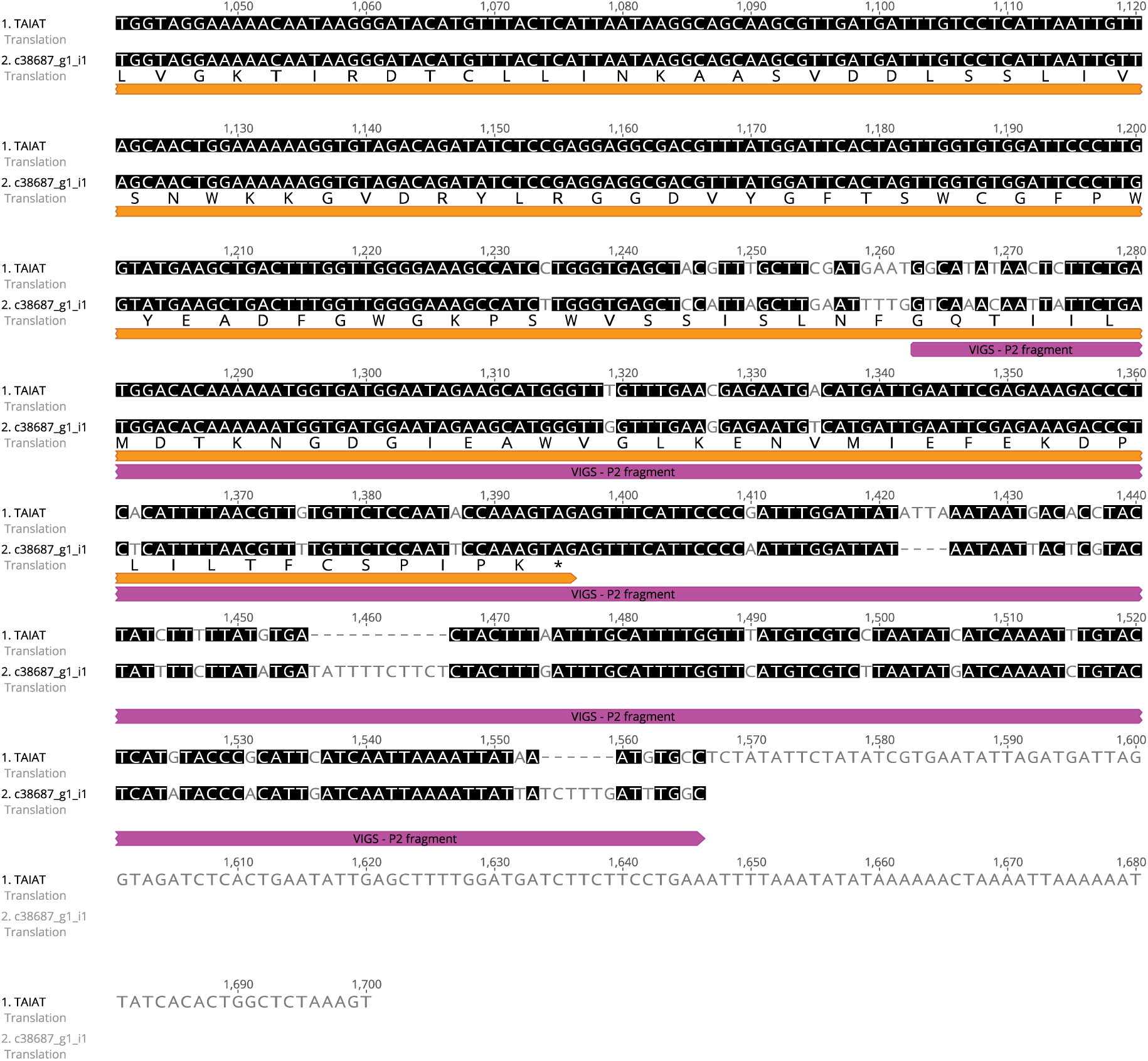
Sequence alignment of TAIAT and ASAT4H transcripts from RNAseq data. Alignments were performed in Geneious V10 using default settings. Conserved nucleotide sequences are colored black, while coding sequencer is denoted by the orange bar. The transcript region targeted by VIGS fragments are annotated by the green (ASAT4H P1) and pink (ASAT4H P2) bars. The c38687_g1_i1 CDS is annotated by an orange bar and amino acid sequence is shown. Gaps are signified by hyphens.

**Figure S7.**
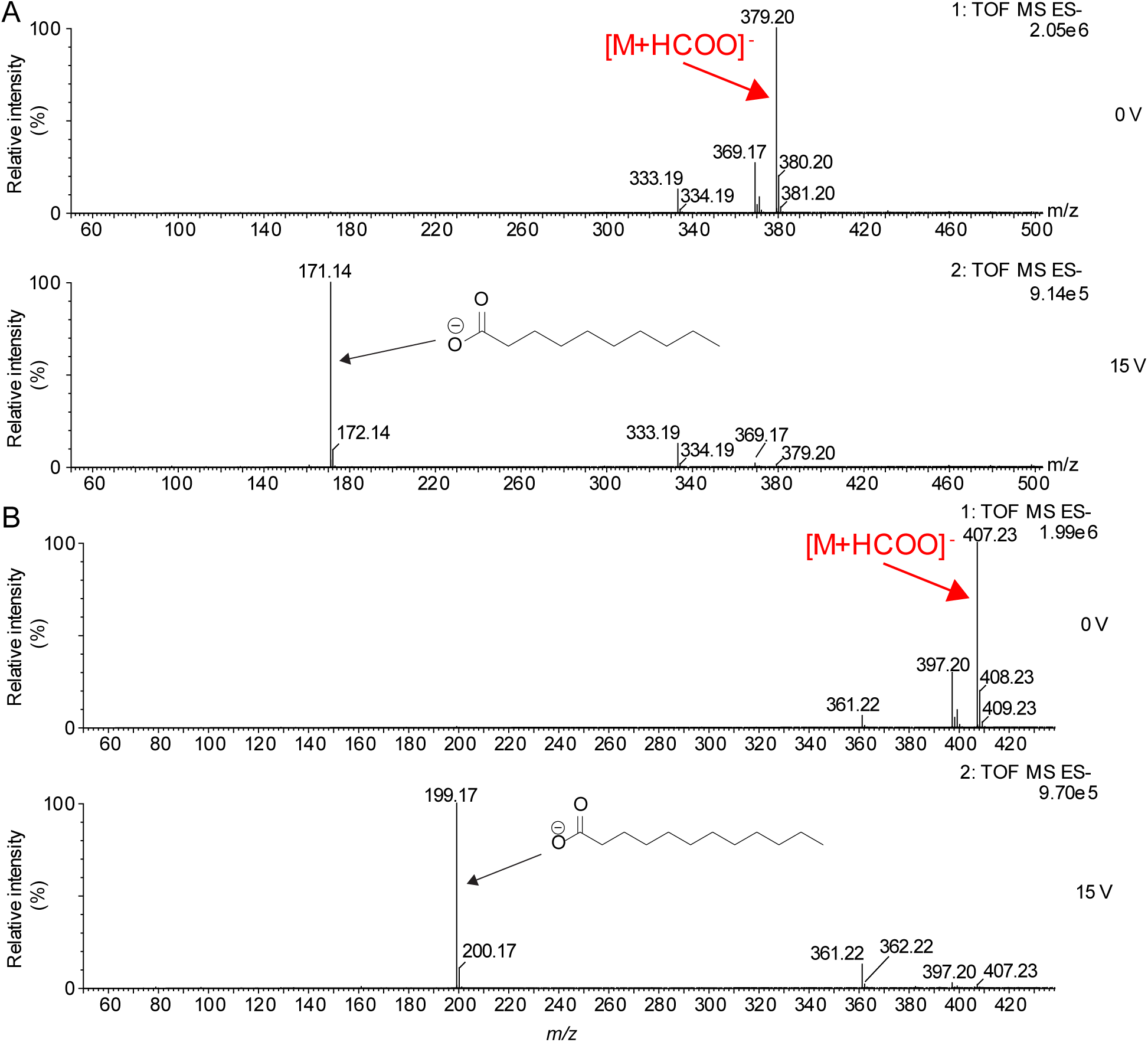
Collision-induced dissociation negative-ion mass spectra of nC10 and nC12 mono-acylated *myo*-inositols generated *in vitro*. A, Collision-induced dissociation of formate adduct of I1:10 yields nC10 carboxylate ion (*m/z* 171.14), as shown in the spectra of pH 6 product (0 V collision potential (top) and 15 V collision potential (bottom)). B, Collision-induced dissociation of I1:12 formate adduct yields nC12 carboxylate ion, as shown in the spectra (0 V collision potential (top) and 15 V collision potential (bottom)).

**Figure S8.**
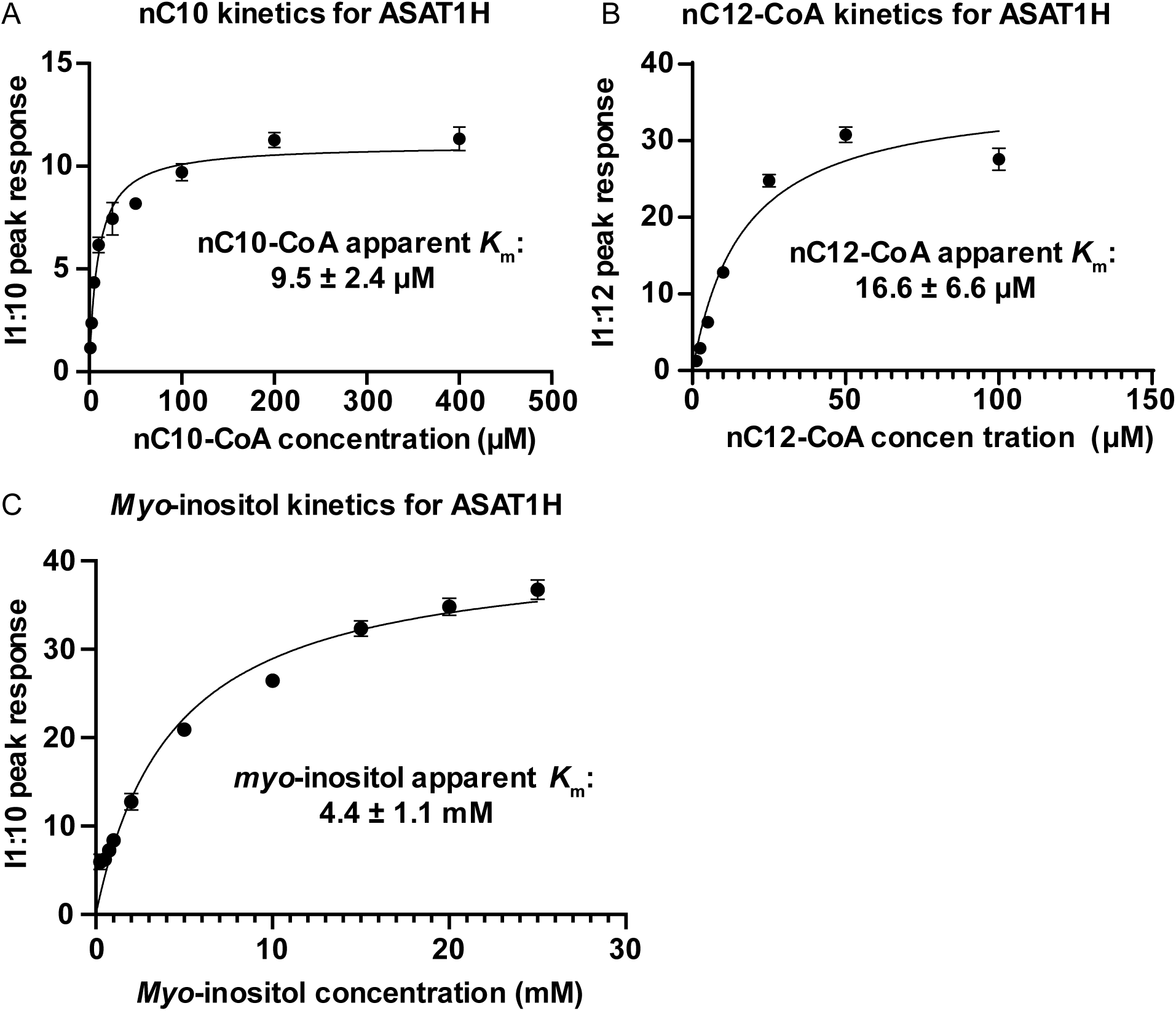
ASAT1H kinetic analysis. Plots for apparent *K*_m_ measurement of A, nC10-CoA, B, nC12-CoA, and C, *myo*-inositol. Apparent *K*_m_ values were calculated using the non-linear regression model in Graphpad prism 8. Products were measured by peak area normalized to internal standard and plotted for the corresponding concentration. Each concentration was performed in triplicate technical replicates with the standard error shown.

**Figure S9.**
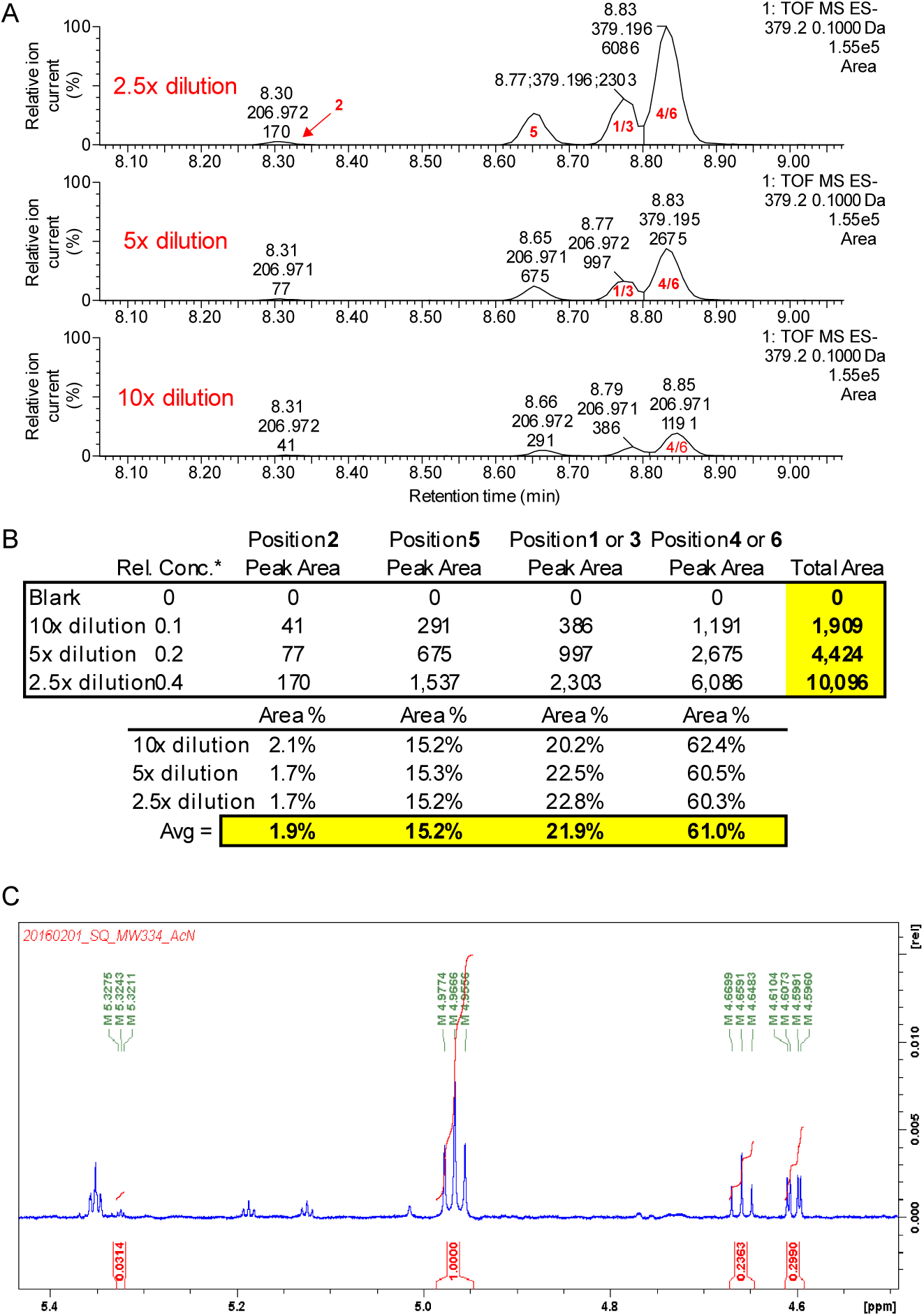
LC-MS and ^1^H NMR analysis of *in vitro* monoacylinositol isomers. A, LC-MS extracted ion chromatograms [M+HCOO]^-^ (*m/z* 379.2) for I1:10 solution of isomers characterized by NMR, and using serial dilutions of (2.5-, 5-, and 10-fold) of the NMR sample. Isomeric ring positions of C10 acylation for chromatographic peaks are labeled in red. B, Integration of peak area from LC-MS analysis of serially diluted samples compared to blank samples. The percentage of total I1:10 peak area is also indicated. C, ^1^H-NMR spectrum of ring hydrogen region for MW 334; C10 mono-acylated *myo*-inositol depicts 4 distinguishable isomers (see Table S6 for summary of study).

**Figure S10.**
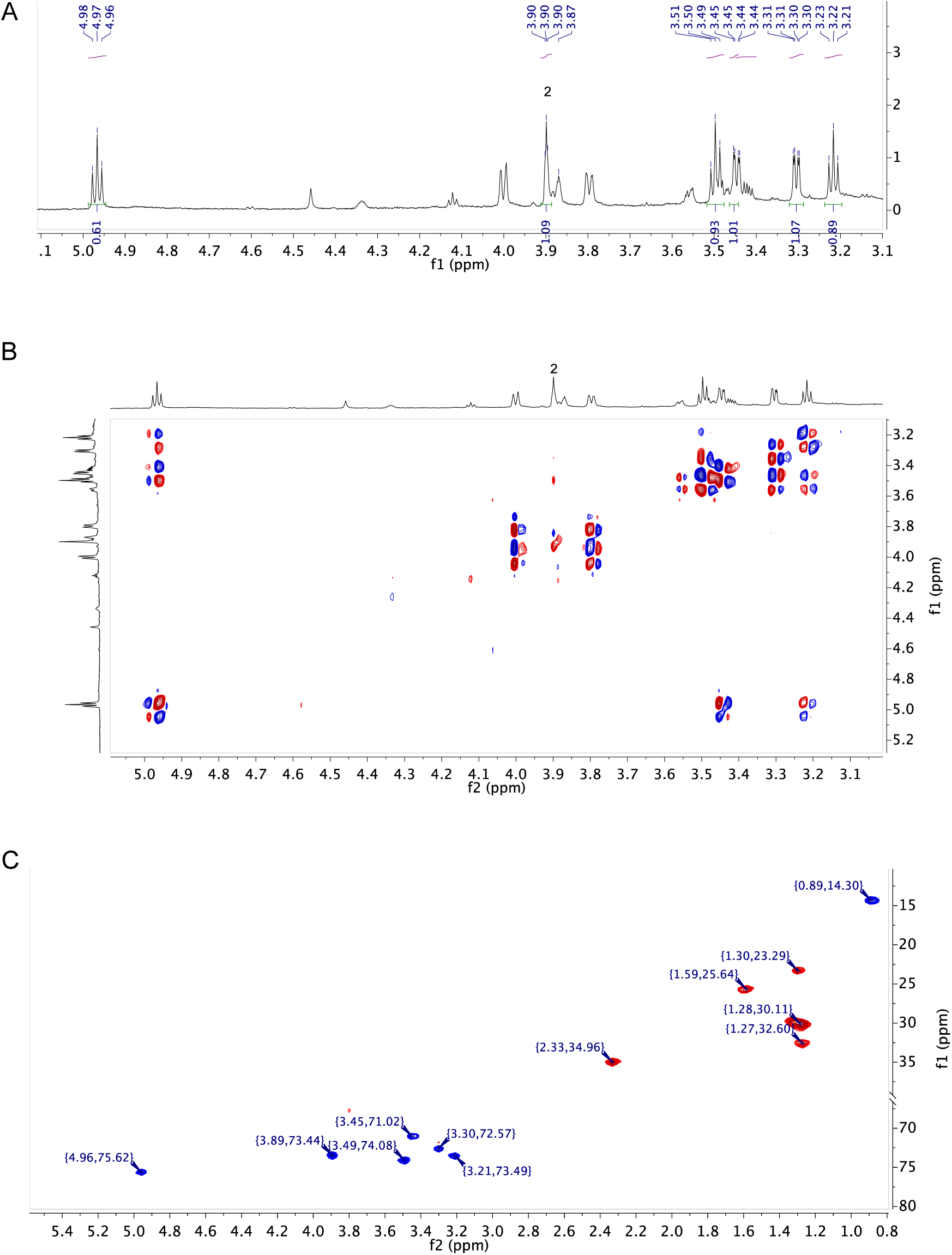
NMR analysis of purified monoacylinositol isomer. A. ^1^H-NMR spectrum of ring hydrogen region for MW 334; purified C10 mono-acylated *myo*-inositol 4/6 isomer; B. ^1^H-^1^H COSY-NMR spectrum of ring hydrogen region for MW 334; purified C10 mono-acylated *myo*-inositol 4/6 isomer; C. HSQC spectrum of for MW 334; purified C10 mono-acylated *myo*-inositol 4/6 isomer.

**Figure S11.**
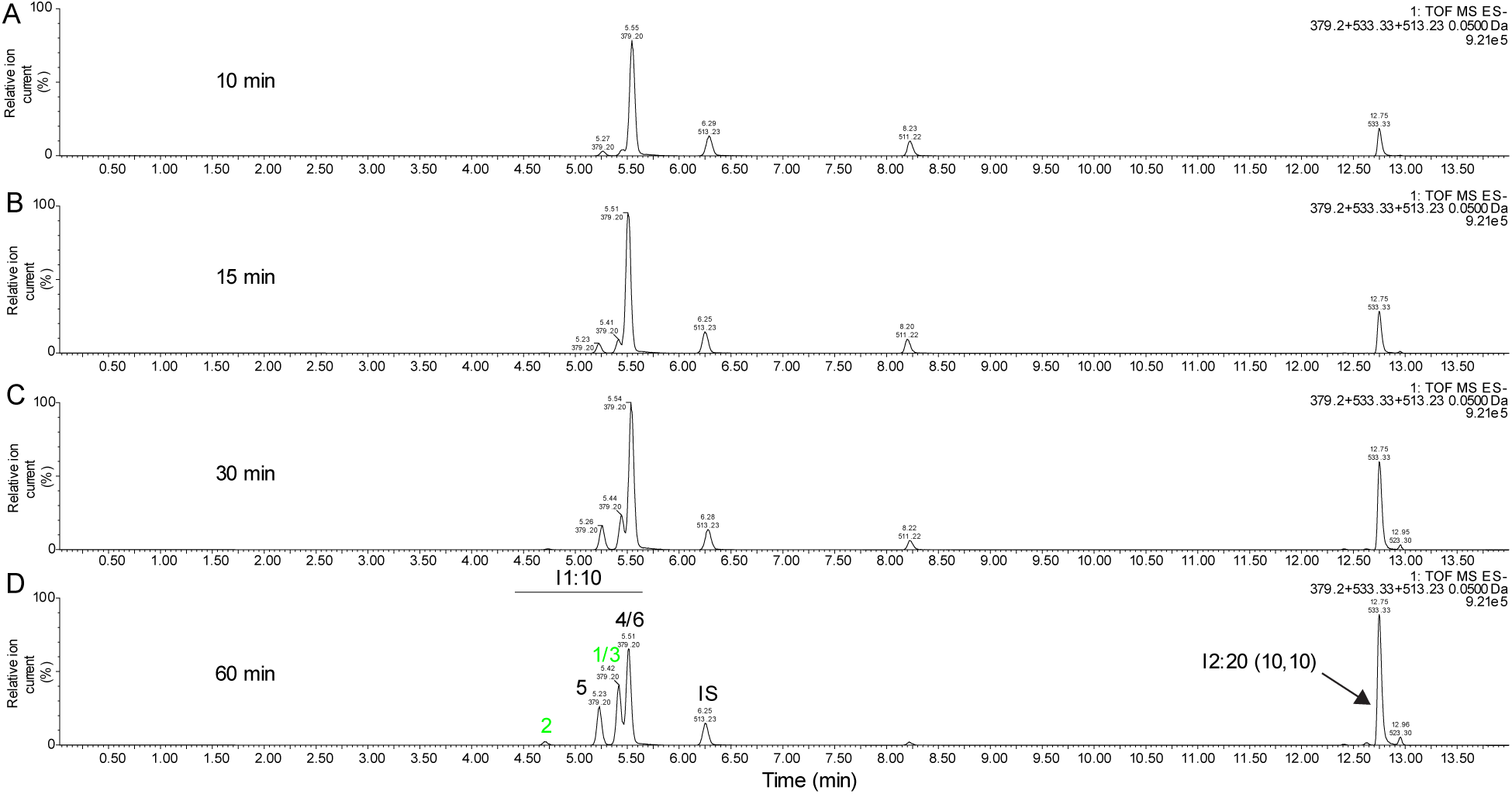
ASAT1H *in vitro* assays at pH 8 reveal a complex mixture of products. LC-MS analysis of ASAT1H *in vitro* assays reactions run at pH 8 for A, 10 minutes, B, 15 minutes, C, 30 minutes, D, 60 minutes. Enzyme was incubated with nC10-CoA and *myo*-inositol. The *m/z* values of the compounds included in this figure: telmisartan (internal standard), *m/z* 513.23; I1:10, *m/z* 379.20; I2:20 (10,10), *m/z* 533.33 with a mass window of 0.05 Da. The samples were run on 14-minute I1:10 method. I1:10 isomers are labeled and *in vivo* positions of medium acyl chains are green.

**Figure S12.**
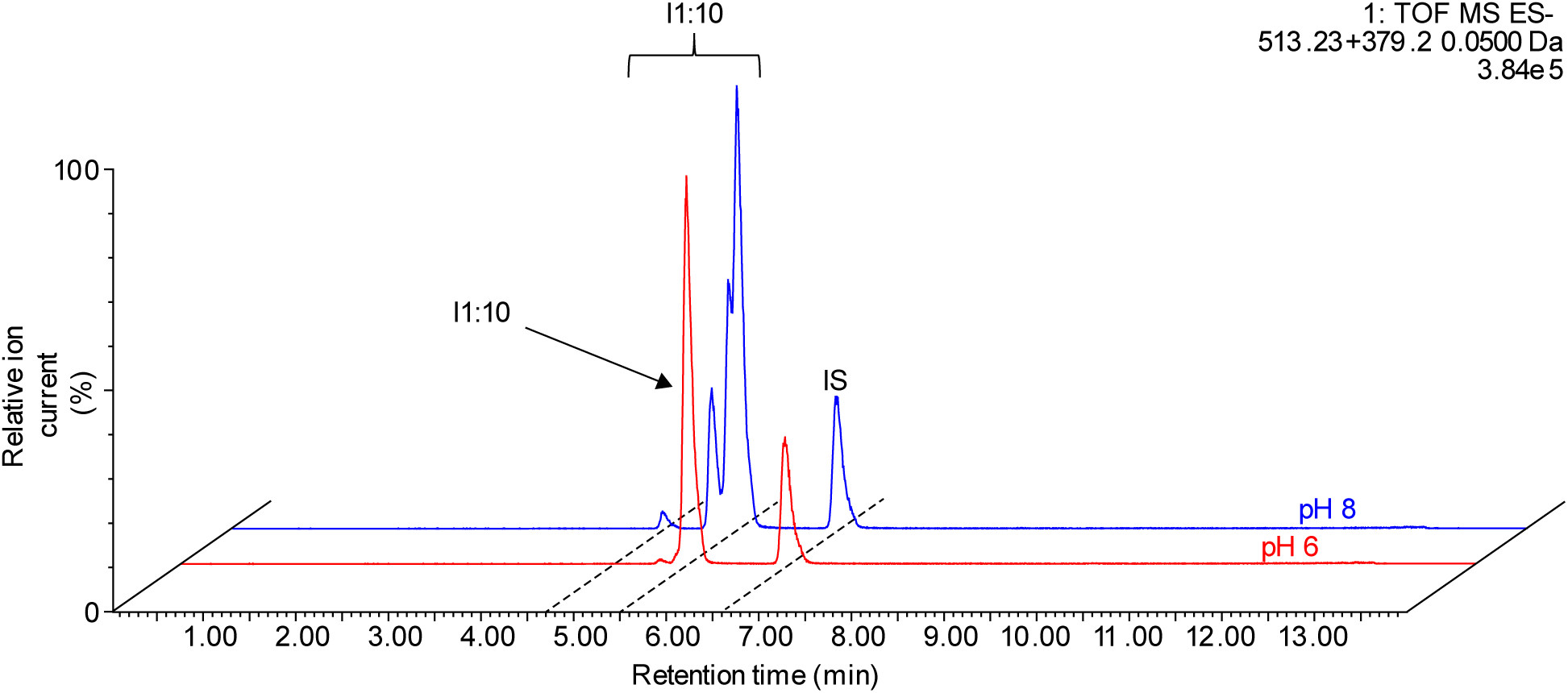
Analysis of pH impact on non-enzymatic rearrangement of ASAT1H I1:10 product. LC-MS analysis of ASAT1H *in vitro* monoacylated product from 60 minute reaction at pH 6 (red) or pH 8 (blue). The *m/z* values of the compounds in the combined extracted ion chromatograms included in this figure: telmisartan (internal standard), *m/z* 513.23; I1:10, *m/z* 379.20 with a mass window of 0.05 Da. Samples were run using a 14-minute chromatography method.

**Figure S13.**
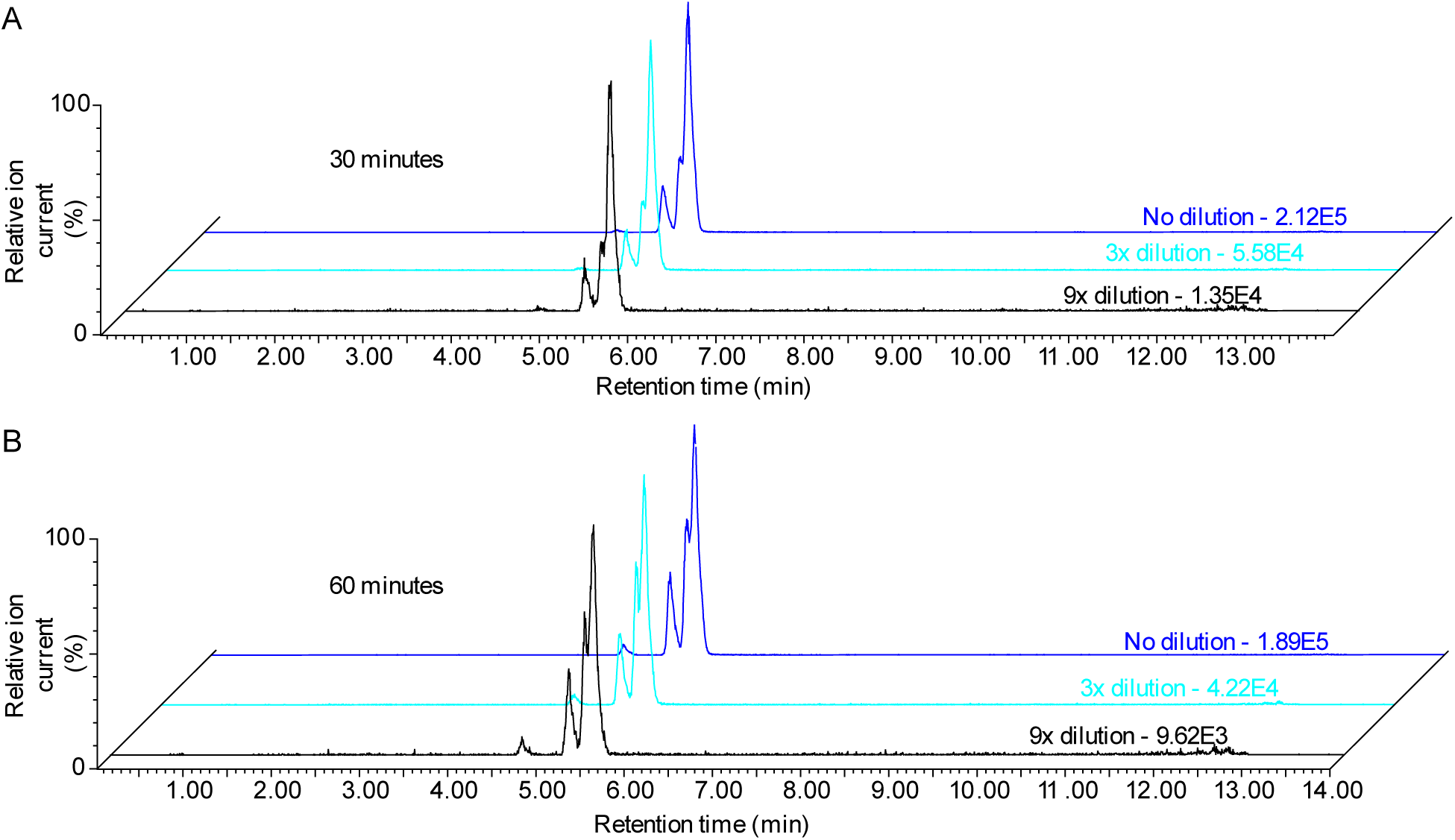
Analysis of dilution impacts on non-enzymatic rearrangement of I1:10 at pH 8. LC-MS analysis of rearrangement products of I1:10 derived from ASAT1H enzyme assays produced at pH 6 and diluted into pH 8 using three input substrate concentrations (1x, 3x, and 9x dilutions). The I1:10 product was purified and added to pH 8 NaPO4 buffer and incubated for 1 hour. The extracted ion chromatograms show the rearrangement of I1:10 at pH 8 for A, 30 minutes and B, 60 minutes.

**Figure S14.**
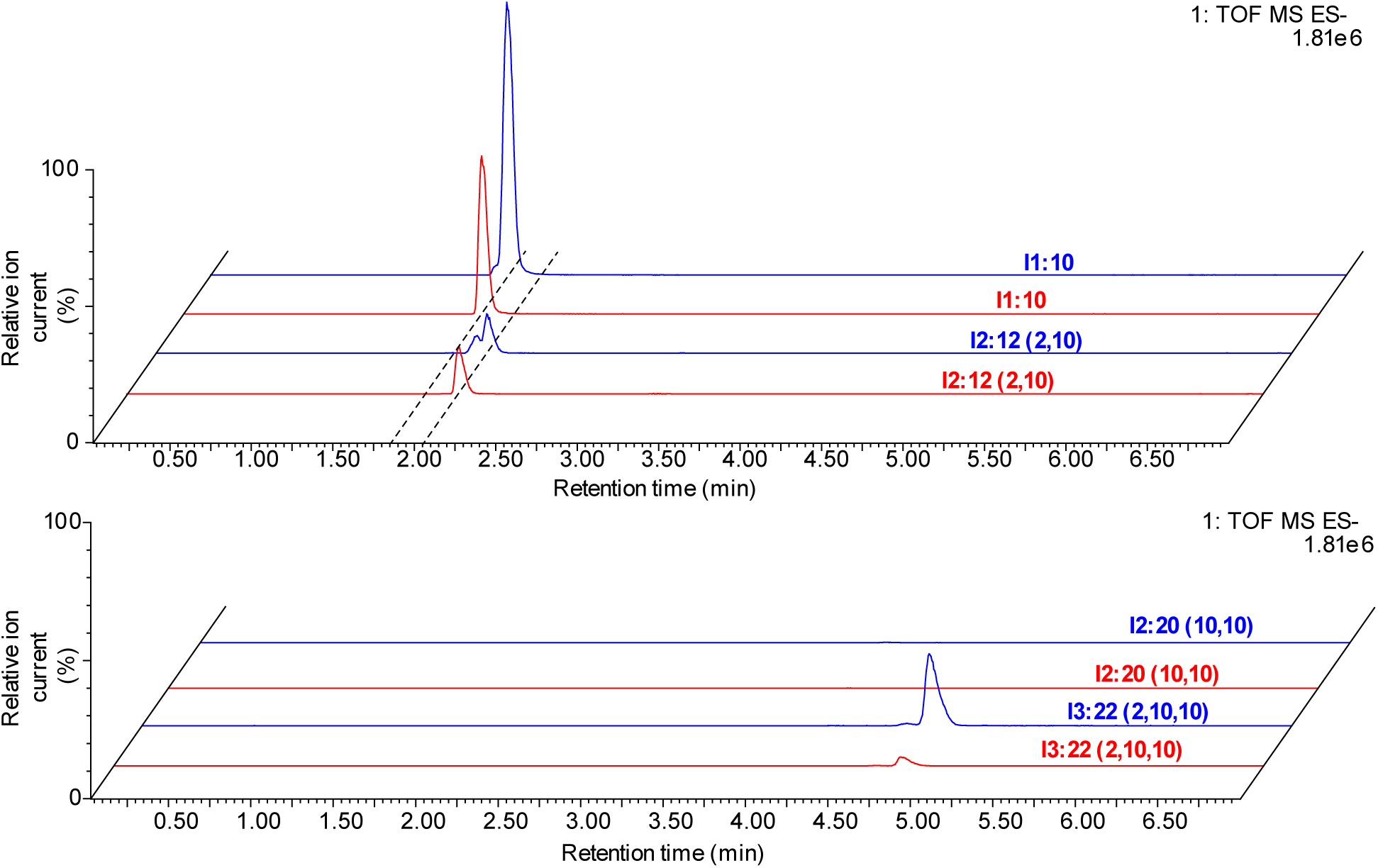
Combined ASAT1H, ASAT3H, and ASAT4H *in vitro* assays. LC-MS analysis of single tube ASAT1H, ASAT3H, and ASAT4H *in vitro* enzyme assays with *myo*-inositol, nC10-CoA and C2-CoA at pH 6 (red) or 8 (blue). Extracted ion chromatograms for I1:10 and I2:12 (2, 10) (top), or I2:20 (10, 10) and I3:22 (2, 10, 10) (bottom) are shown. The *m/z* values of the compounds in the combined extracted ion chromatograms included in this figure: I1:10, *m/z* 379.20; I2:12 (2,10), *m/z* 421.20; I2:20 (10,10), *m/z* 533.33; I3:22 (2,10,10), *m/z* 575.35 with a mass window of 0.05 Da.

**Figure S15.**
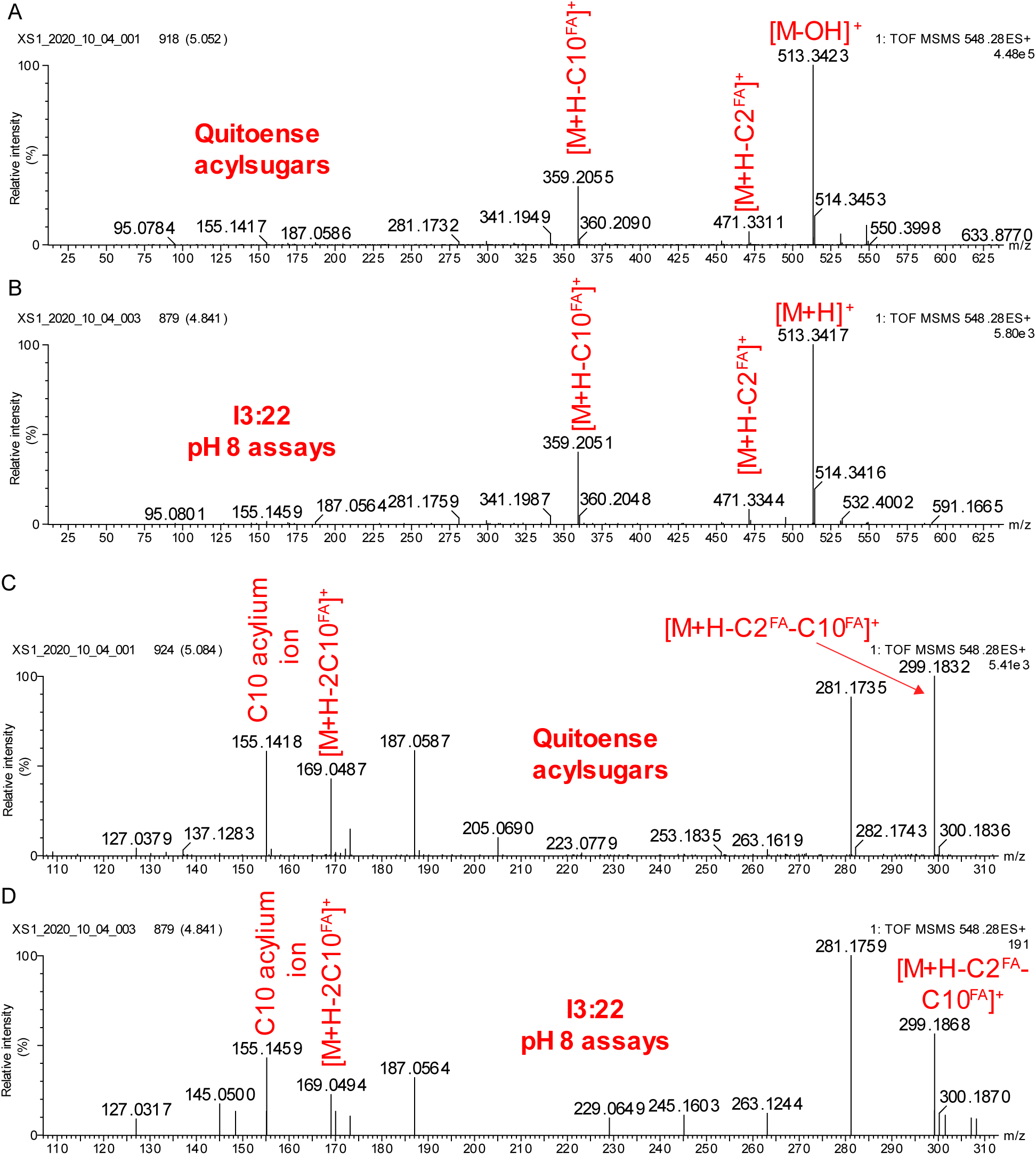
ASAT1H, ASAT3H, and ASAT4H enzyme assay product and *S. quitoense in vivo* acylsugar positive-ion mode LC-MS/MS spectra. Fragmentation of A, *S. quitoense* I3:22 (2,10,10) and B, *in vitro* pH 8 I3:22 product yields product ions that indicate the neutral loss of C2 and C10 carboxylic acids. C-D, Subsection (*m/z* 110-310) of spectra from Panel A and Panel B that contains an ion with the neutral loss of 2 C10 acyl chains. Key product ions that show neutral losses of acyl groups are annotated with superscript FA indicating neutral losses of carboxylic acids (plus ammonia) from [M+NH_4_]^+^. C10 acylium ions (RCO^+^) are annotated for product ions at *m/z* 155.1. Fragmentation methods are detailed in Materials and Methods.

**Figure S16.**
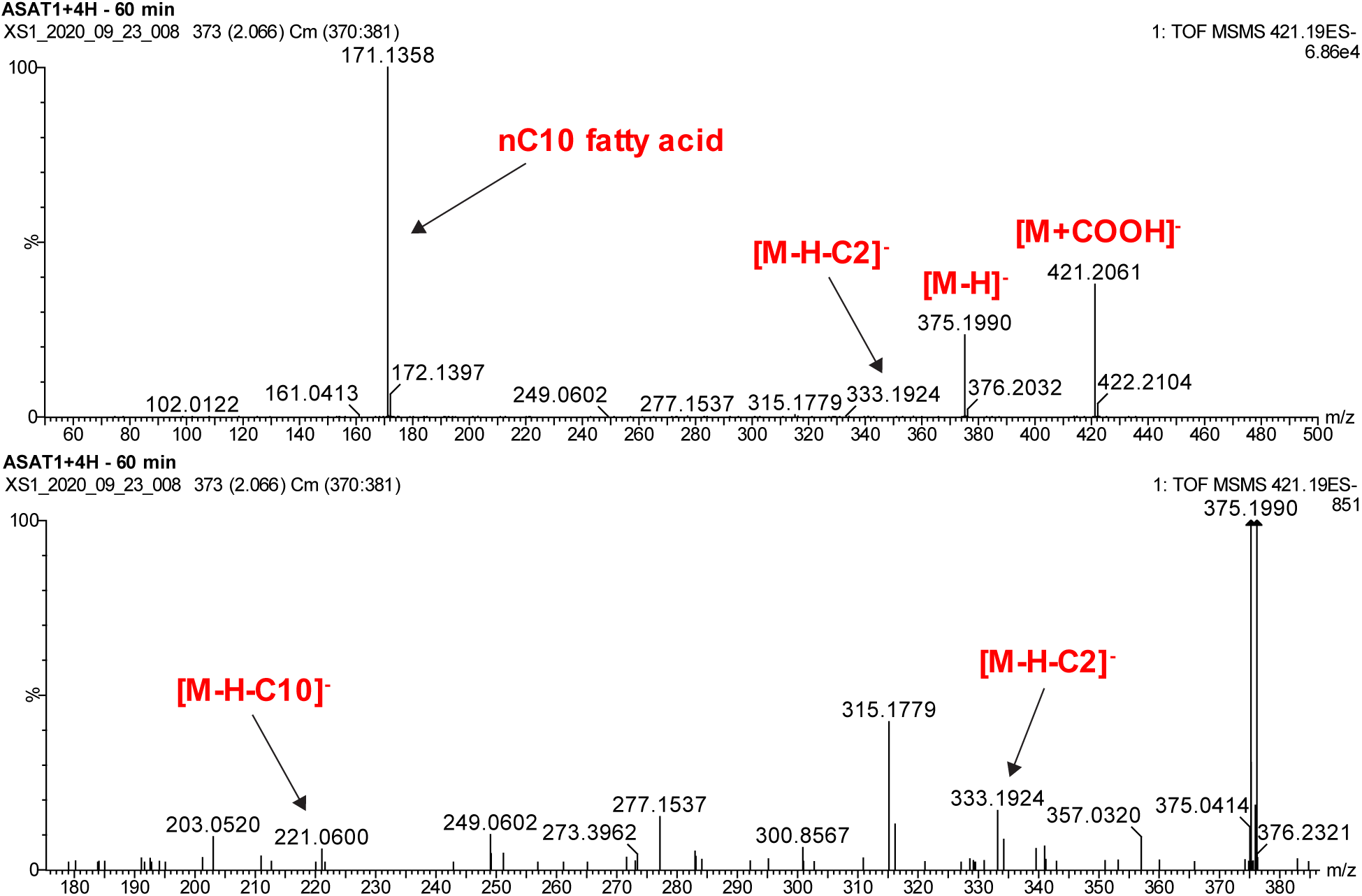
LC-MS/MS spectra of ASAT1 and ASAT4H assay products. I2:12 (2,10) produces a C10 carboxylate (*m/z* 171.14). Fragmentation also produces ions from ketene neutral loss of C10 or C2 acyl chains. The bottom spectrum shows an expanded and magnified view for the spectrum in the mass range *m/z* 175 to 387.

**Figure S17.**
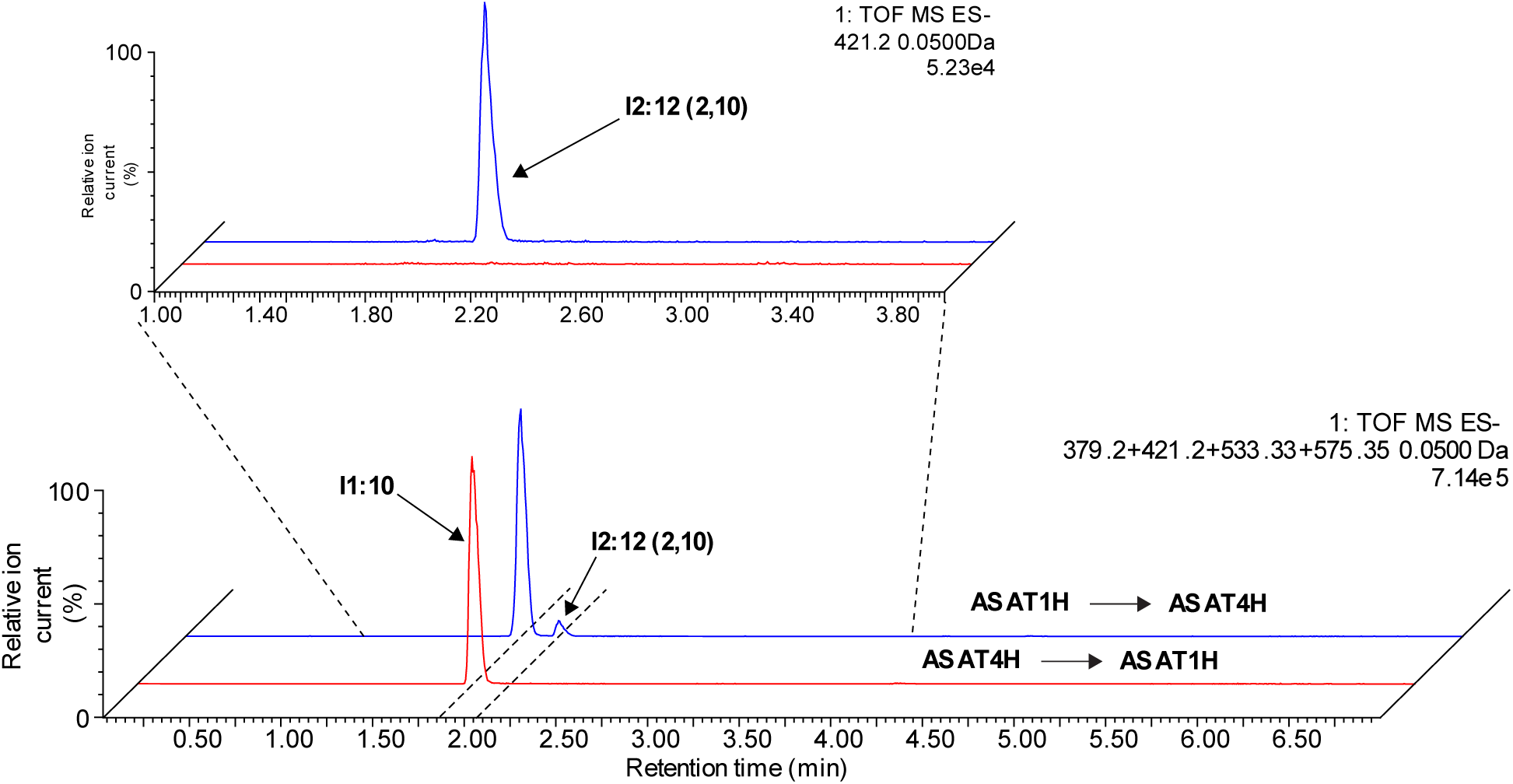
Sequential ASAT1H and ASAT4H *in vitro* assays. LC-MS analysis of ASAT1H and ASAT4H sequential *in vitro* enzyme assays with *myo*-inositol, nC10-CoA and C2-CoA substrates. Reactions with ASAT4H as the first enzyme, heat inactivation, then ASAT1H (red) and with ASAT1H as the first enzyme, heat inactivation, then ASAT4H (blue) are shown. The *m/z* values of the compounds in the combined extracted ion chromatograms included in this figure: I1:10, *m/z* 379.20; I2:12 (2,10), *m/z* 421.20; I2:20 (10,10),b *m/z* 533.33; I3:22 (2,10,10), *m/z* 575.35 with a mass window of 0.05 Da. I2:12 (2,10) extracted ion chromatogram from min 1-4 shows no I2:12 in reaction with ASAT4H followed by ASAT1H.

**Figure S18.**
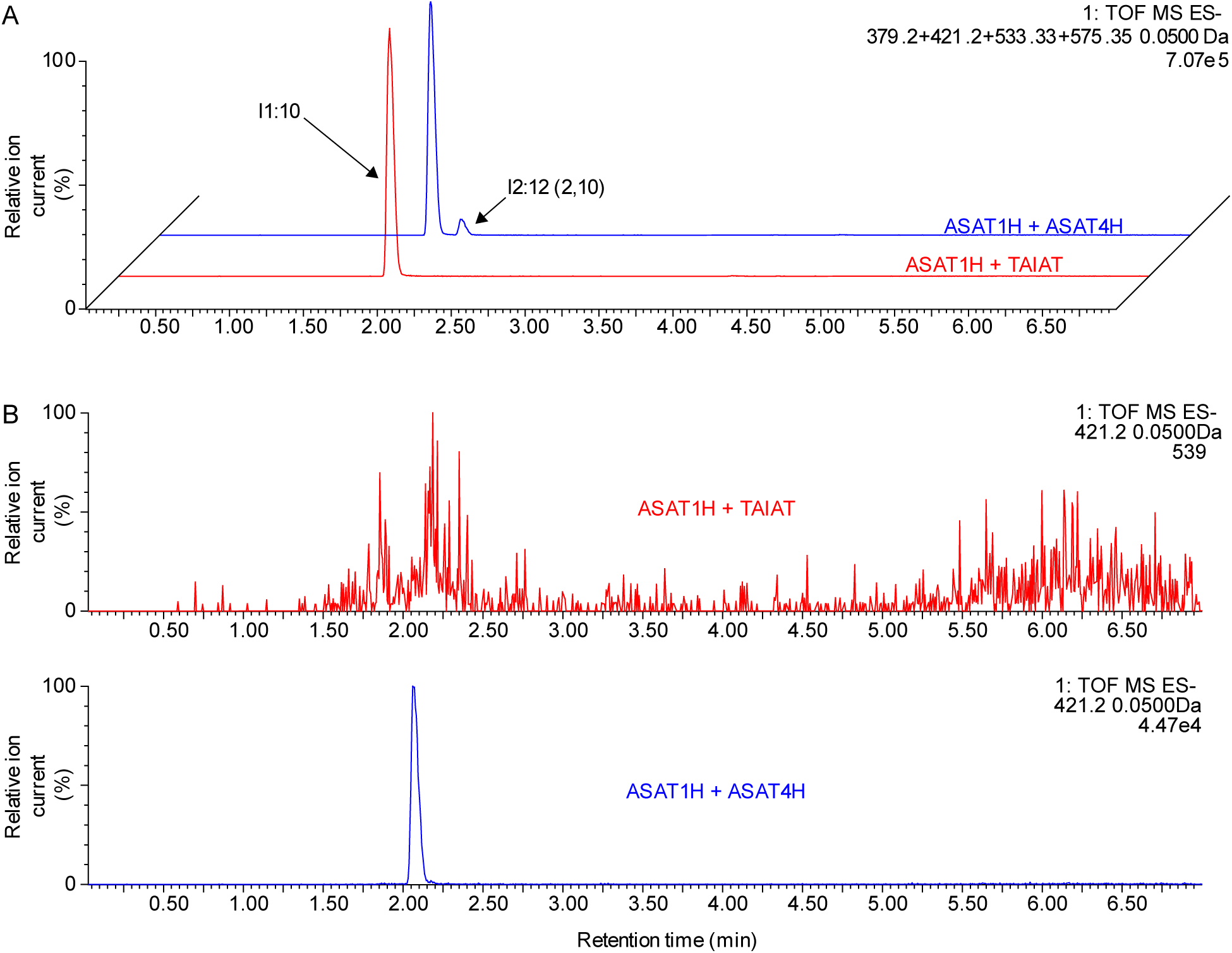
ASAT1H and ASAT4H or TAIAT combined *in vitro* assays. A, LC-MS analysis of ASAT1H and ASAT4H (red) or TAIAT (blue) combined *in vitro* enzyme assays using *myo*-inositol, nC10-CoA and C2-CoA and run for 30 minutes. B, Extracted ion chromatograms of I2:12 (2,10) from ASAT1H and TAIAT (top) or ASAT4H (bottom) reactions are shown for clarity and comparison. Note that the TAIAT reaction product MS sensitivity is set approximately 100-fold higher. The *m/z* values of the compounds in the combined extracted ion chromatograms included in this figure: I1:10, *m/z* 379.20; I2:12 (2,10), *m/z* 421.20; I2:20 (10,10), *m/z* 533.33; I3:22 (2,10,10), *m/z* 575.35, each with a mass window of 0.05 Da.

**Figure S19.**
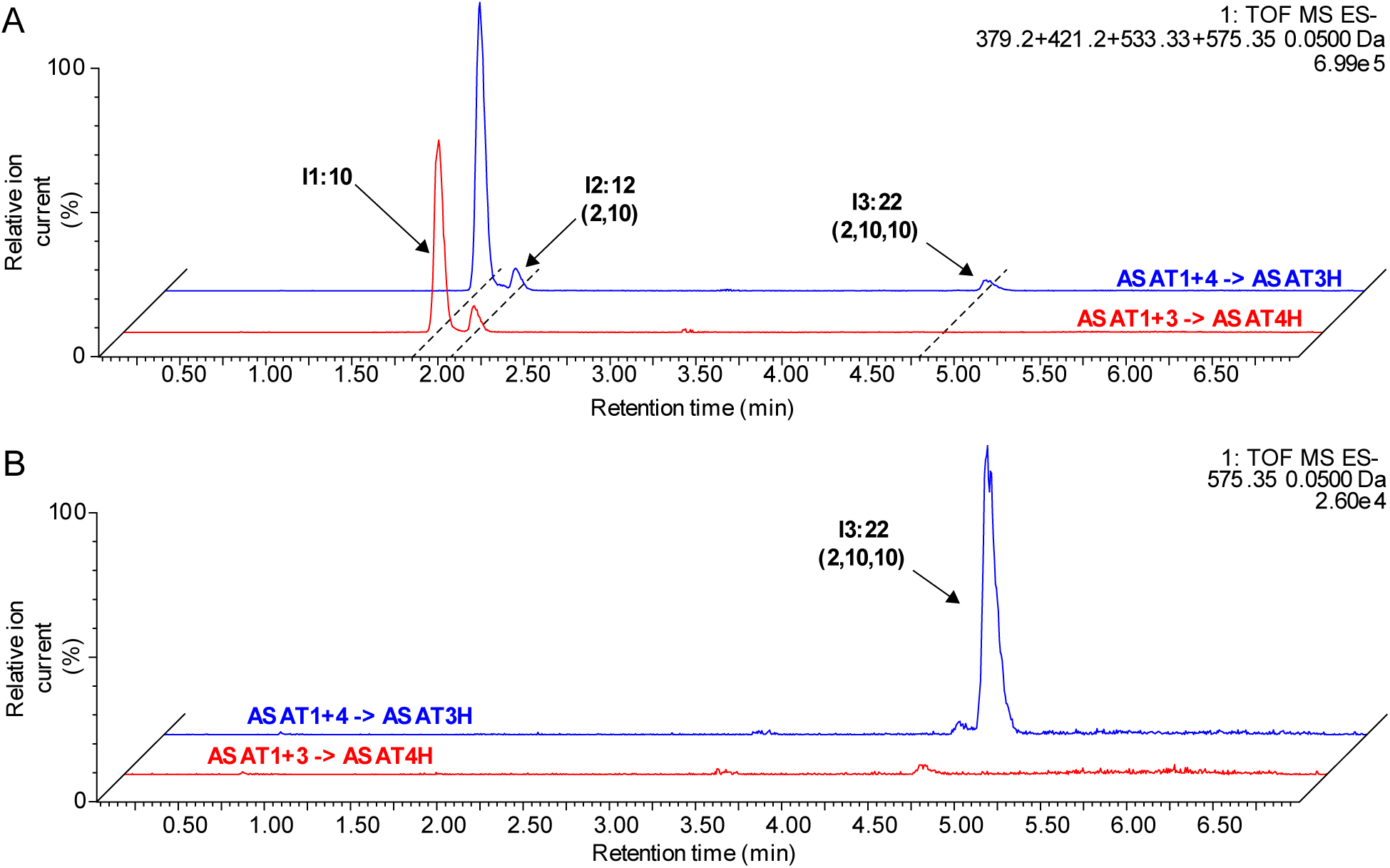
ASAT1H, ASAT3H, and ASAT4H sequential *in vitro* assays. A, LC-MS analysis of ASAT1H, ASAT3H, and ASAT4H *in vitro* sequential enzyme assays with *myo*-inositol, nC10-CoA and C2-CoA. Sequential reactions with ASAT1H and ASAT3H followed by heat inactivation, then ASAT4H (red) or ASAT1H and ASAT4H followed by heat inactivation, then ASAT3H (blue) are shown. B, An extracted ion chromatogram including I3:22 (2,10,10) is shown for clarity and comparison. The *m/z* values of the compounds in the combined extracted ion chromatograms included in this figure: I1:10, *m/z* 379.20; I2:12 (2,10), *m/z* 421.20; I2:20 (10,10), *m/z* 533.33; I3:22 (2,10,10), *m/z* 575.35, all with a mass window of 0.05 Da.

Table S1. Acylsugar analysis of *S. quitoense* ASAT3H and ASAT4H-targeted VIGS plant samples.

Acylsugars were measured as described in Materials and Methods. Acylsugar response is acylsugar peak area divided by internal standard peak area. The acylsugar response is divided by leaf dry weight to give the leaf dry weight normalized response.

Table S2. Acylsugar analysis of *S. quitoense* ASAT1H-targeted plants.

Acylsugars were measured as described in Materials and Methods. Acylsugar response is acylsugar peak area divided by internal standard peak area. The acylsugar response is divided by leaf dry weight to give the leaf dry weight normalized response.

Table S3. Percent identity of *S. quitoense* ASAT homolog protein sequence to SlASATs.

Table S4. Measurements of *K*m values of nC10-CoA, nC12-CoA, and *myo*-inositol for ASAT1H.

In each case, one substrate was held constant at a saturating concentration, while the other was varied in concentration. Enzyme velocity was measured as a function of the I1:10 or I1:12 peak area normalized to the internal standard. Samples were analyzed using LC-MS-MS using multiple reaction monitoring on a Waters Acquity TQD instrument. nC10-CoA was used as the acyl-CoA for the *myo*-inositol kinetic measurements.

Table S5. NMR chemical shifts for product of ASAT1H at pH 6.

Table S6. Characterization of I1:10 isomer mixture by LC-MS and ^1^H NMR.

Table S7. Oligonucleotide sequences used in this study.

## Materials and Methods

### Candidate identification and phylogenetic characterization

Identification of suitable gene candidates was performed in our previous work (Leong et al., 2020). In short, sequences were selected based on the presence of the HXXXD motif (0 mismatches) and the DFGWG motif (1 mismatch allowed) using the ‘search for motif’ function in Geneious R.8.1.9. Those sequences that qualified were filtered to those between 400-500 amino acids, and by relative position of the two motifs (D’Auria, 2006). Eggplant putative ASAT homologs were obtained by blasting Sl-ASAT1-4 and Solyc07g043670 coding sequence against the Eggplant genome v3 CDS sequences on Sol Genomics Network using blastn.

Sequences were aligned against other characterized BAHD sequences and characterized ASATs using MEGA X (Kumar et al., 2018). The evolutionary history was inferred by using the Maximum Likelihood method and Whelan And Goldman + Freq. model (Whelan and Goldman, 2001). The bootstrap consensus tree inferred from 1000 replicates (Felsenstein, 1985) is taken to represent the evolutionary history of the taxa analyzed (Felsenstein, 1985). Branches corresponding to partitions reproduced in less than 50% bootstrap replicates are collapsed. Initial tree(s) for the heuristic search were obtained automatically by applying Neighbor-Join and BioNJ algorithms to a matrix of pairwise distances estimated using the JTT model, and then selecting the topology with superior log likelihood value. A discrete Gamma distribution was used to model evolutionary rate differences among sites (5 categories (+*G*, parameter = 1.9258)). This analysis involved 63 amino acid sequences. All positions with less than 30% site coverage were eliminated, i.e., fewer than 70% alignment gaps, missing data, and ambiguous bases were allowed at any position (partial deletion option). There were a total of 459 positions in the final dataset.

### Heterologous protein expression and purification from *Escherichia coli*

Heterologous protein expression was achieved using pET28b(+) (EMD Millipore, Burlington, MA), in which open reading frames for the enzymes were cloned into using doubly digested vectors of either BamHI/XhoI (ASAT4H), NheI/XhoI (ASAT1H), or NheI/NotI (c8981_g1_i1). The doubly digested vectors were assembled with a single fragment containing the ORF containing 5′ and 3′ adapters for Gibson assembly using 2x NEB Hifi Mastermix (NEB, Ipswich, MA). The finished constructs were transformed into BL21 Rosetta (DE3) cells (EMD Millipore, Burlington, MA) and verified using colony PCR and Sanger sequenced using T7 promoter and terminator primers. Oligonucleotide sequences are presented in Table S7.

LB overnight cultures with kanamycin (50 µg/mL) and chloramphenicol (33 µg/mL) were inoculated with a single colony of the bacterial strain containing the desired construct and incubated at 37 °C, 225 rpm, overnight. Larger cultures were inoculated 500:1 with the same antibiotics and incubated at the same temperature and speed. OD600 of the cultures was monitored until between 0.5 and 0.8. Cultures were chilled on ice for 15 minutes, at which IPTG was added to a final concentration of 50 µM for all BAHD sequences except for IAT, which was incubated with 300 µM IPTG. Cultures were incubated at 16 °C, 180 rpm for 16 hours. Note: all of the following steps were processed on ice.

Cultures were centrifuged at 4,000*g* for 10 minutes to collect the cells and repeated until all the culture was processed (4 °C). The cell pellets were resuspended in 25 mL of extraction buffer (50 mM NaPO_4_, 300 mM NaCl, 20 mM imidazole, 5 mM 2-mercaptoethanol, pH 8.0) by vortexing. The cell suspension was sonicated for 8 cycles (30 seconds on, intensity 4, 30 seconds on ice). The cellular extracts were centrifuged at 30,000*g* for 10 minutes. The supernatant was transferred into another tube and centrifuged again at the same speed and duration. Ni-NTA resin (Qiagen, Hilden, Germany) was centrifuged at 1,000*g* for 1 minute, resuspended in 1 mL of extraction buffer. The slurry was centrifuged again at 1,000*g* for 1 minute and the supernatant was decanted. The resin was resuspended using the crude extract and incubated at 4 °C, nutating for 1 hour. The slurry was centrifuged at 3,200*g* for 5 minutes, and supernatant decanted. The resin was resuspended in 5 mL of extraction buffer and transferred to a gravity flow column (Biorad, Hercules, CA). After loading, the resin was washed with 3 column volumes of extraction buffer (∼30 mL). The resin was further washed with 1 column volume of wash buffer (extraction buffer with 40 mM imidazole). The remaining protein was eluted and collected using 2 mL of elution buffer after a 1 minute incubation with the resin. The elution was diluted into 15 mL of storage buffer (extraction buffer, but no imidazole). This elution was concentrated using 30 kDa centrifugal filter units (EMD Millipore, Burlington, MA), and repeated until diluted 1,000 fold. An equal volume of 80% glycerol was added to the elution, mixed, and stored at -20 °C.

### General enzyme assays

Assays were run in 100 mM sodium phosphate, pH 6 or pH 8 at a total volume of 60 µL with pH 6 as the default unless otherwise stated. Acyl-CoAs were added to a final concentration of 100 µM. Non-acylated acceptors were added at a final concentration of 1 mM. 6 µL of enzyme was added to each reaction. The assays were incubated at 30 °C for 30 minutes unless otherwise stated. After the incubation, 2 volumes of stop solution—composed of 1:1 (v/v) of acetonitrile and isopropanol with 0.1% formic acid and 1 µM telmisartan as internal standard (Sigma-Aldrich, St. Louis, MO)—were added to the assays and mixed by pipetting. Reactions were stored in the -20 °C freezer for 20 minutes and centrifuged at 17,000*g* for 5 minutes. The supernatant was transferred to LC-MS tubes and stored at -20 °C.

ASAT1H timepoint assays at pH 8 were scaled up to 200 µL reactions with the same proportions of reagents to allow time-point aliquots. Assays that tested ASAT1H and ASAT4H together were run with 50 mM *myo*-inositol, 3 µL of each enzyme was added to each 60 µL reaction, and incubated for 30 minutes, unless otherwise stated. Sequential assays testing ASAT1H or 4H order included an initial incubation of 30°C for 30 minutes with the first enzyme (volume is 60 µL at this point), a 70°C incubation for 10 minutes to inactivate the enzyme, followed by addition of 3 µL each of 2 mM decanoyl-CoA, acetyl-CoA, and 3 µL of the second enzyme with a 30 minute incubation at 30°C. Joint assays with ASAT1, 3, and 4H together included 2 µL of each enzyme, 50 mM *myo*-inositol, and were incubated for 2 hours. Sequential assays testing ASAT1H, 3H, and 4H included a similar format: 2 µL of ASAT1H and ASAT3 or 4H (60 µL total volume) with a 1 hour incubation at 30°C, assays were incubated at 70°C for 10 minutes to inactivate enzymes, followed by the addition of 2 µL of the last enzyme and 3 µL each of 2 mM decanoyl and acetyl-CoA and a 1 hour incubation at 30°C.

For kinetic analysis, conditions were used such that the enzyme amount and reaction time were in the linear range. For each substrate (nC10, nC12, and *myo*-inositol), the other substrate was held at saturating concentrations. The reactions were run for 20 minutes, performed in triplicate, and stopped by the addition of 2 volumes of stop solution. Samples were analyzed as described in LC-MS section. Nonlinear regression was performed using standard Michaelis-Menten kinetics model in Graphpad Prism 8 (Graphpad Software).

For rearrangement assays, a 10 mL reaction with 1 mL of ASAT1H in 100 mM sodium phosphate, pH 6 with 100 µM decanoyl-CoA and 75 mM *myo*-inositol was run for 3 hours. Reactions were stopped with 2 volumes of stop solution (1:1 acetonitrile:isopropanol with 0.1% formic acid), dried down by speedvac and resuspended in 8 mL of water:acetonitrile (6:4) and purified by method for the single product NMR analysis in **NMR analysis of monoacylinositols** below except that the purified fraction was resuspended in 2 mL of water:acetonitrile with 0.1% formic acid and dried down before storage at -20°C. Assays were run in 100 mM sodium phosphate, pH 6 and 8. I1:10 product was resuspended in an ethanol:water mixture (1:1) and 1/60^th^ of a volume was added to the reaction. Timepoint assays were run and sampled at 5, 10, 15, 30, 45, and 60 minutes. Serial dilutions of monoacylinositol products confirmed that proportions of isomers are maintained by LC-MS (Figure S9, Table S5, and Table S6)

### Mono-acylated enzyme assays for NMR analysis

Assays for single product NMR analyses were run in 100 mM ammonium acetate, pH 6.0 at a total volume of 60 mL. nC10-CoA and *myo*-inositol were added to a final concentration of 400 µM and 30 mM, respectively. 6 mL of enzyme solution was added to the reactions purified from 6 L of *E.coli* culture. Reactions were incubated at 30 °C for 3 hours. After incubation, 2 volumes of stop solution—1:1 acetonitrile:isopropanol containing 0.1% formic acid—were added to the assays and mixed by pipetting. The multiproduct NMR analysis was similar except it was run in 50 mM ammonium acetate, pH 6 with 100 µM nC10-CoA and 75 mM *myo*-inositol, and incubated at 30°C for 45 minutes. Reactions were evaporated to dryness using the speedvac. In both cases, the products were purified using a Waters 2795 Separations module equipped with LKB Bromma 2211 Superrac fraction collector with automated fraction collection.

### Purification and NMR analysis of monoacylinositols

**For the single mono-acylated NMR product analysis:** the residue was resuspended in a 6 mL mixture of water and acetonitrile (6:4) with 0.1% formic acid by vortexing and transferred to LC-MS vials for semipreparative purification. Samples were purified on the semipreparative LC using 200 µL injections at a flow rate of 1.5 mL/minute using a C18 semipreparative column at 30 °C (Acclaim C18 5 µm 120 Å, 4.6 x 150 mm, ThermoFisher). Solvents A and B were: water with 0.1% formic acid, and acetonitrile, respectively. The 24 minute linear LC gradient was as follows: 5% B at 0 minutes; 25% B at 1.00 minute; 35% B at 20.00 minutes; 100% B at 21.00 minutes; 100% B until 22.00 minutes; 5% B at 22.01 minutes; 5% B until 24.00 minutes. The duration of collection of each fraction was 15 seconds. The 14-minute I1-10 method described in the Methods section was used to analyze fractions using mass spectrometry. Fraction 72 was evaporated to dryness in a speedvac. The residue was resuspended in a 1 mL mixture of water:acetonitrile (6:4) with 0.1% formic acid. The solution was dried in a speedvac and resuspended in deuterated acetonitrile before being transferred to Shigemi tubes for NMR analysis. ^1^H, ^13^C, gCOSY, gHSQC, gHMBC and NOESY NMR experiments were performed at the Max T. Rogers NMR Facility at Michigan State University using a Bruker Avance 900 spectrometer equipped with a TCI triple resonance probe. All spectra were referenced to non-deuterated CD_3_CN solvent signals (δ_H_ = 1.94 and δ_C_ = 1.32, 118.26 ppm). The ^1^H spectra were recorded at 900 MHz, while the ^13^C spectra were recorded at 225 MHz.

**For the NMR sample containing multiple monoacylinositol isomers:** the dried acylsugar was resuspended in a 6 mL mixture of water and acetonitrile (6:4) with 0.1% formic acid by vortexing and the solvent was evaporated under vacuum using a Thermo Savant SPD 131 DDA Speedvac concentrator with BOC Edwards XDS scroll pump. The residue was reconstituted in acetonitrile:isopropanol (ACN:IPA) (1:1) with sonication, combined to a single 18 x 150 mm tube and concentrated to dryness under vacuum using the Speedvac. Approximately 300 µL of ACN:IPA (1:1) was added to the tube, sonicated, and centrifuged. The supernatant was then transferred to an LC autosampler vial with limited volume insert for semi-preparative HPLC purification (∼250 µL sample volume). The purification was performed by semipreparative LC via two 200 µL injections using a C18 semipreparative column at 50 °C (Acclaim C18 5 µm 120 Å, 4.6×150 mm). To maximize sample recovery, after the first injection, approximately 150 µL of AcN:IPA was added to the culture tube, sonicated and centrifuged. The supernatant was then transferred to the same LC autosampler vial with limited volume insert for a second injection.

One minute fractions were collected in Pyrex glass culture tubes (18 × 150 mm, total volume 3 mL each fraction) and labeled by minute. The 50-minute LC gradient had solvents A: 0.15% formic Acid in water; B: acetonitrile; and column wash with C: dichloromethane: acetone: methanol (v/v/v, 1;1:1). The linear gradient began with a hold at 5% B from 0-1 minute, ramp from 5-20% B from 1-2 minutes, ramp from 20-40% B from 2-25 minutes The fractions were combined according to the observed purity and the abundance of the ion at *m/z* 379.20, consistent with [M+HCOO]^-^ of C10 monoacylinositols. Fractions 16-21 had the greatest abundance. Those fractions were combined and concentrated to dryness under vacuum for NMR, and ramp from 40-100% B from 25-26 minutes. The solvent profile continued with a hold at 100% B from 26-28 minutes, ramp from 100% B to 100% C from 28-29 minutes, hold at 100% C from 29-39 minutes, and ramp from 100% C to 100% B from 39-40 minutes. The final stages of the solvent gradient were a ramp from 100-5% B from 40-41 minutes, and hold at 5% B for 41-50 minutes. R analysis. The sample was dissolved in ∼250 µL acetonitrile-*d*_3_ with vortex mixing and transferred to a solvent matched Shigemi tube and analyzed by ^1^H-NMR. The NMR spectrum was recorded using a Bruker Avance 900 MHz NMR spectrometer equipped with a TCI triple-resonance inverse detection cryoprobe at the Max T. Rogers NMR facility at Michigan State University.

For the LC-MS analysis of monoacylinositol isomers (Figure S9), the following LC gradient used solvents A: 10 mM ammonium formate, pH 2.8 in water; and B: acetonitrile. Samples were analyzed on a Waters Acquity UPLC coupled to a Waters Xevo G2-XS QToF mass spectrometer with an Acquity UPLC HSS-T3 C18 column (2.1 mm x 100 mm x 1.8 µm). The gradient with a flow rate of 0.5 mL/minutes is as follows: Hold at 99% A from 0-1 minute, ramp to 50% A from 1-5 minutes, ramp to 100% A from 10-10.01 minutes, hold at 100% A from 10.01-12.50 minutes, ramp to 99% A from 12.50-12.51 minutes, and hold at 99% A from 12.51-15.00 minutes.

### LC-MS analysis

LC-MS samples (both enzyme assays and plant samples) were analyzed on a Waters Acquity UPLC coupled to a Waters Xevo G2-XS QToF mass spectrometer (Waters Corporation, Milford, MA). Sample injection volume was 10 µL. LC-MS methods were performed with gradient elution using an Ascentis Express C18 HPLC column (10 cm x 2.1 mm, 2.7 µm) (Sigma-Aldrich, St. Louis, MO) at 40 °C using the following solvents: 10 mM ammonium formate, pH 2.8 as solvent A, and 100% acetonitrile as solvent B. A flow rate of 0.3 mL/minute was used unless otherwise specified. All LC-MS was performed with electrospray ionization in either positive or negative ion mode as described.

A 7-minute linear elution gradient consisted of 5% B at 0 minutes, 60% B at 1 minute, 100% B at 5 minutes, held at 100% B until 6 minutes, 5% B at 6.01 minutes and held at 5% until 7 minutes.

A 14 minute I1-10 linear elution gradient consisted of 5% B at 0 minutes; 25% B at 1 minute; 50% B at 10 minutes; 100% B at 12 minutes; 5% B at 12.01 minutes, 5% B at 14.00 minutes. This method is the default 14-minute method unless otherwise stated.

An alternative 14-minute linear elution gradient was used for the comparison of I3:22 metabolites extracted from *S. quitoense* or generated in an enzyme assay containing ASAT1H and TAIAT or ASAT4H. The 14-minute linear elution gradient consisted of 5% B at 0 minutes; 60% B at 1 minute; 100% B at 12 minutes; 100% B at 13 minutes; 5% B at 13.01 minutes, 5% B at 14.00 minutes.

A 21-minute linear elution gradient of 5% B at 0 minutes, 60% B at 3 minutes, 100% B at 15 minutes, held at 100% B until 18 minutes, 5% B at 18.01 minutes and held at 5% until 21 minutes.

For ESI-MS settings: capillary voltage, 2.00 kV; source temperature, 100 °C; desolvation temperature, 350 °C; desolvation nitrogen gas flow rate, 600 liters/hour; cone voltage, 40 V; mass range, *m/z* 50-1000 (with spectra accumulated at 0.1s per function). Three quasi-simultaneous acquisition functions were used to acquire spectra at different collision potentials (0, 15, and 35 V). Lock mass correction was performed using leucine enkephalin as the reference for data acquisition.

For ESI+ MS settings: capillary voltage, 3.00 kV; source temperature, 100 °C; desolvation temperature, 350 °C; desolvation nitrogen gas flow rate, 600 liters/hour; cone voltage, 35 V; mass range, *m/z* 50-1000 (with spectra accumulated at 0.1 s per function). Two acquisition functions were used to acquire spectra at different collision potential settings (0, 10-60 V). Lock mass correction was performed using leucine enkephalin as the reference for data acquisition.

LC-MS/MS analyses were performed same instruments and columns as other LC-MS analysis. Ion source parameters were as described above. Survey scans were acquired over *m/z* 50 to 1,000 with 0.3 s per scan. Collision potential was ramped from 10-60 V. For I2:12, the formate adduct (*m/z* 421.20) was used. For I3:22, the ammonium adduct (*m/z* 548.38) and formate adduct (*m/z* 575.34) were used for analysis.

For *K*_m_ measurements, the same column, 7 minute LC method, and solvents were used for the analysis on the Waters Acquity TQD triple quadrupole mass spectrometer coupled to a Waters Acquity UPLC. The parameters used for the mass spectrometer are as follows: Capillary voltage, 2.5 kV; Cone voltage, 30 V; Source temperature, 130°C; Desolvation temperature, 350°C; Cone gas flow, 20 L/hr, Desolvation gas flow, 800 L/hr. The mass pairs used are as follows: Telmisartan, *m/z* 513 > 287; I1:10, *m/z* 379 > 171; *m/z* I1:12, 407 > 199.

### VIGS analysis

pTRV2-LIC was digested using PstI-HF to generate the linearized vector. The linearized vector was purified using a 1% agarose gel and gel extracted using an Omega EZNA gel extraction kit. Fragments were amplified using PCR with adapters for ligation into pTRV2-LIC. Both the PCR fragment and the linearized vector were incubated in separate 5 µL reactions using NEB 2.1 as buffer with T4 DNA polymerase and 5 mM dATP or dTTP (PCR insert/Vector). The reactions were incubated at 22 °C for 30 minutes, subsequently incubated at 70 °C for 20 minutes. The reactions were then stored on ice. 1 µL of the pTRV2-LIC reaction and 2 µL of the PCR-LIC reaction were mixed by pipetting. Reactions were incubated at 65 °C for 2 minutes, then 22 °C for 10 minutes. After which the constructs were transformed into chemically competent *E.coli* cells.

Constructs were tested for the presence of the insert using colony PCR and pTRV2-LIC-seq-F/R primers showing a 300 bp insertion. Positive constructs were miniprepped (Qiagen, Hilden, Germany) and sanger sequenced using the same primers. Sequenced constructs and pTRV1 were transformed into agrobacterium strain, GV3101, using the protocol described previously except on LB plates with kanamycin (50 µg/mL), rifampicin (50 µg/mL), and gentamycin (10 µg/mL). Colonies were assayed for the presence of the insert using the colony PCR and the pTRV2-LIC-seq-F/R primers previously described. The presence of the pTRV1 vector in GV3101 was assayed using colony PCR primers, pTRV1-F/R.

Protocol adapted from: (Velásquez et al., 2009). Seeds were germinated using an incubation in 10% bleach for 30 minutes, followed by 5-6 washes with water. Seeds were transferred to a petri dish with Whatman paper and water in the bottom of the dish. Seeds were stored in a lab drawer until hypocotyls emerge, at which point they were moved to a window sill. Once, cotyledons had emerged, seedlings were transferred to peat pots and grown for approximately 1 week under 16/8 day/night cycle at 24 °C. At 2 days pre-inoculation, LB cultures (Kan/Rif/Gent) were inoculated with the cultures used for leaf inoculation. The strains had constructs containing the gene of interest (GOI) in pTRV2-LIC, an empty vector pTRV2-LIC, and pTRV1. Cultures were grown overnight at 30 °C with shaking at 225 rpm. Larger cultures composed of induction media (4.88 g MES, 2.5 g glucose, 0.12 g sodium phosphate monobasic monohydrate in 500 mL, pH 5.6, 200 µM acetosyringone), were inoculated using a 25:1 dilution of the overnight culture (50 mL total). The larger culture was incubated at 30 °C, 225 rpm, overnight. Cells were harvested by centrifugation at 3,200*g* for 10 minutes. Cell pellets were resuspended in 1 volume of 10 mM MES, pH 5.6, 10 mM MgCl_2_. Cells were gently vortexed to resuspend the pellet. Cell suspensions were centrifuged at 3,200*g* for 10 minutes. Cell pellets were resuspended in 10 mL of 10 mM MES, pH 5.6, 10 mM MgCl_2_. The OD600 values were measured for each of the cultures. Cell suspensions were diluted using the same buffer to an OD600 of 1. Acetosyringone was added to the pTRV1 cell suspension to a final concentration of 400 µM. The different pTRV2-LIC constructs were mixed into 50 mL conical tubes with an equal volume of pTRV1 suspension, resulting in a final acetosyringone concentration of 200 µM. Individual seedlings were inoculated through the abaxial side of the cotyledon. Plants were incubated at 22 °C and shaded for 24 hours. After 24 hours, the plants were returned to 16/8h day-night cycles at the same temperature. Approximately 3 weeks later, the plants were sampled for acylsugars and RNA using a bisected leaf for each experiment.

**Note:** Inoculation timing is very important; the cotyledons should be inoculated after they have expanded, but before the first two true leaves have fully emerged.

### Acylsugar analysis

The interactive protocol for acylsugar extracts is available at Protocols.io at: https://dx.doi.org/10.17504/protocols.io.xj2fkqe

The acylsugar extraction protocol was described in (Leong et al., 2019). LC-MS conditions used for acylsugar analysis were described the LC-MS analysis section.

### qRT-PCR analysis

RNA was extracted with the RNeasy Plant Mini Kit including on-column DNase digestion (Qiagen), according to the manufacturer’s instructions. RNA was quantified with a Nanodrop 2000c instrument (Thermo Fisher Scientific). cDNA was synthesized using 1 mg of the isolated RNA and SuperScript II Reverse Transcriptase (Invitrogen). The cDNA samples were diluted 20-fold (10-fold initial dilution and 2-fold dilution before qPCRs). qPCRs (10 uL) were created with SYBR Green PCR Master Mix with 1 µL of cDNA added (Thermo Fisher Scientific). Primers were used at a final concentration of 200 nM. RT_ASAT1H_F/R, RT_ASAT3H_F/R, RT_ASAT4H_F/R, RT_TAIAT_F/R,RT_Actin_F/R, and RT_EF1a_F/R primers were used to detect ASAT1H, ASAT3H, ASAT4H, TAIAT, ACTIN, and EF1a transcripts, respectively (Supplemental Table S7). Reactions were carried out with a QuantStudio 7 Flex Real-Time PCR System (Applied Bio-systems) by the Michigan State University RTSF Genomics Core. The following temperature cycling conditions were applied: 50°C for 2 min, 95°C for 10 min, and 40 cycles of 95°C for 15 s and 60°C for 1 min. Relative expressions of ASAT1H, ASAT3H, ASAT4H, and TAIAT were calculated with the DDCt method (Pfaffl, 2001) and normalized to the geometric mean of ACTIN and EF1a transcript levels. The mean expression values of the transcripts in the control plants were used for normalization. Three to four technical replicates were used for all the qPCRs.

### Accession numbers

ASAT homolog sequence data are available in GenBank as follows: TAIAT, MT024677; c38687_g1_i1, MT024678; c38687_g2_i1, MT024679; c39979_g2_i1, ON005014; c8981_g1_i1, ON005015.

## References

Ahuja I, Kissen R, Bones AM (2012) Phytoalexins in defense against pathogens. Trends in Plant Science 17: 73–90

Burke B, Goldsby G, Mudd JB (1987) Polar epicuticular lipids of *Lycopersicon pennellii*. Phytochemistry 26: 2567–2571

D’Auria JC (2006) Acyltransferases in plants: a good time to be BAHD. Current Opinion in Plant Biology 9: 331–340

Fan P, Leong BJ, Last RL (2019) Tip of the trichome: evolution of acylsugar metabolic diversity in Solanaceae. Current Opinion in Plant Biology 49: 8–16

Fan P, Miller AM, Liu X, Jones AD, Last RL (2017) Evolution of a flipped pathway creates metabolic innovation in tomato trichomes through BAHD enzyme promiscuity. Nature Communications 8: 2080

Fan P, Miller AM, Schilmiller AL, Liu X, Ofner I, Jones AD, Zamir D, Last RL (2016) *In vitro* reconstruction and analysis of evolutionary variation of the tomato acylsucrose metabolic network. Proceedings of the National Academy of Sciences 113: E239–48

Fan P, Wang P, Lou Y-R, Leong BJ, Moore BM, Schenck CA, Combs R, Cao P, Brandizzi F, Shiu S-H, et al (2020) Evolution of a plant gene cluster in Solanaceae and emergence of metabolic diversity. eLife 9: 2020.03.04.977231

Felsenstein J (1985) Confidence limits on phylogenies: an approach using the bootstrap. Evolution 39: 783–791

Ghosh B, Westbrook TC, Jones AD (2014) Comparative structural profiling of trichome specialized metabolites in tomato (*Solanum lycopersicum*) and *S. habrochaites*: acylsugar profiles revealed by UHPLC/MS and NMR. Metabolomics 10: 496–507

Herrera-Salgado Y, Garduño-Ramírez ML, Vázquez L, Rios MY, Alvarez L (2005) *Myo*- inositol-derived glycolipids with anti-inflammatory activity from *Solanum lanceolatum*. Journal of Natural Products 68: 1031–1036

Howe GA, Jander G (2008) Plant immunity to insect herbivores. Annu. Rev. Plant Biol 59: 41– 66

Hurney SM (2018). Strategies for profiling and discovery of acylsugar specialized metabolites. [Doctoral dissertation, Michigan State University]. Proquest.

Kim J, Kang K, Gonzales-Vigil E, Shi F, Jones AD, Barry CS, Last RL (2012) Striking natural diversity in glandular trichome acylsugar composition is shaped by variation at the Acyltransferase2 locus in the wild tomato *Solanum habrochaites*. Plant Physiology 160: 1854–70

King RR, Calhoun LA (1988) 2,3-Di-*O*- and 1,2,3-tri-*O*-acylated glucose esters from the glandular trichomes of *Datura metel*. Phytochemistry 27: 3761–3763

King RR, Pelletier Y, Singh RP, Calhoun LA (1986) 3,4-Di-*O*-isobutyryl-6-*O*-caprylsucrose: the major component of a novel sucrose ester complex from the type B glandular trichomes of *Solanum berthaultii* hawkes (Pl 473340). J Chem Soc, Chem Commun 1078–1079

Kumar S, Stecher G, Li M, Knyaz C, Tamura K (2018) MEGA X: molecular evolutionary genetics analysis across computing platforms. Molecular biology and evolution 35: 1547–1549

Landis JB, Miller CM, Broz AK, Bennett AA, Carrasquilla-Garcia N, Cook DR, Last RL, Bedinger PA, Moghe GD (2021) Migration through a major Andean ecogeographic disruption as a driver of genetic and phenotypic diversity in a wild tomato species. Molecular Biology and Evolution 38: 3202–3219

Landry LG, Chapple CCS, Last RL (1995) Arabidopsis mutants lacking phenolic sunscreens exhibit enhanced ultraviolet-B injury and oxidative damage. Plant Physiology 109: 1159–1166

Leong BJ (2019) Analysis of acylglucose and acylinositol biosynthetic mechanisms in Solanaceae species. [Doctoral dissertation, Michigan State University]. Proquest.

Leong BJ, Hurney SM, Fiesel PD, Moghe GD, Jones AD, Last RL (2020) Specialized metabolism in a nonmodel nightshade: trichome acylinositol biosynthesis. Plant Physiology 183: 915–924

Leong BJ, Lybrand DB, Lou Y-R, Fan P, Schilmiller AL, Last RL (2019) Evolution of metabolic novelty: a trichome-expressed invertase creates specialized metabolic diversity in wild tomato. Science Advances 5: eaaw3754

Liu X, Enright M, Barry CS, Jones AD (2017) Profiling, isolation and structure elucidation of specialized acylsucrose metabolites accumulating in trichomes of *Petunia* species. Metabolomics 13: 85

Lou Y-R, Anthony TM, Fiesel PD, Arking RE, Christensen EM, Jones AD, Last RL (2021) It happened again: convergent evolution of acylglucose specialized metabolism in black nightshade and wild tomato. Science Advances 7: eabj8726

Lybrand DB, Anthony TM, Jones AD, Last RL (2020) An integrated analytical approach reveals trichome acylsugar metabolite diversity in the wild tomato *Solanum pennellii*. Metabolites 10: 1–25

Maldonado E, Torres FR, Martínez M, Pérez-Castorena AL (2006) Sucrose esters from the fruits of *Physalis nicandroides* var. attenuata. Journal of Natural Products 69: 1511–1513

Mandal S, Ji W, McKnight TD (2020) Candidate gene networks for acylsugar metabolism and plant defense in wild tomato *Solanum pennellii*. The Plant Cell 32: 81–99

Massalha H, Korenblum E, Tholl D, Aharoni A (2017) Small molecules below-ground: the role of specialized metabolites in the rhizosphere. Plant Journal 90: 788–807

Matsuzaki T, Shinozaki Y, Suhara S, Shigematsu H, Koiwai A (1989) Isolation and characterization of tetra- and triacylglucose from the surface lipids of *Nicotiana miersii*. Agricultural and Biological Chemistry 53: 3343–3345

Mithöfer A, Boland W (2012) Plant defense against herbivores: chemical aspects. Annu Rev Plant Biol 63: 431–450

Moghe GD, Leong BJ, Hurney S, Jones AD, Last RL (2017) Evolutionary routes to biochemical innovation revealed by integrative analysis of a plant-defense related specialized metabolic pathway. eLife 6: 1–33

Mugford ST, Qi X, Bakht S, Hill L, Wegel E, Hughes RK, Papadopoulou K, Melton R, Philo M, Sainsbury F, et al (2009) A serine carboxypeptidase-like acyltransferase is required for synthesis of antimicrobial compounds and disease resistance in oats. The Plant Cell 21: 2473– 2484

Nadakuduti SS, Uebler JB, Liu X, Jones AD, Barry CS (2017) Characterization of trichome-expressed BAHD acyltransferases in *Petunia axillaris* reveals distinct acylsugar assembly mechanisms within the Solanaceae. Plant Physiology 175: 36–50

Ning J, Moghe GD, Leong BJ, Kim J, Ofner I, Wang Z, Adams C, Jones AD, Zamir D, Last RL (2015) A feedback-insensitive isopropylmalate synthase affects acylsugar composition in cultivated and wild tomato. Plant Physiology 169: 1821–1835

Schilmiller AL, Charbonneau AL, Last RL (2012) Identification of a BAHD acetyltransferase that produces protective acyl sugars in tomato trichomes. Proceedings of the National Academy of Sciences 109: 16377–16382

Schilmiller AL, Gilgallon K, Ghosh B, Jones AD, Last RL (2016) Acylsugar acylhydrolases: carboxylesterase catalyzed hydrolysis of acylsugars in tomato trichomes. Plant Physiology 170: 1331–1344

Schilmiller AL, Moghe GD, Fan P, Ghosh B, Ning J, Jones AD, Last RL (2015) Functionally divergent alleles and duplicated loci encoding an acyltransferase contribute to acylsugar metabolite diversity in *Solanum* trichomes. The Plant Cell 27: 1002–1017

Shirley AM, McMichael CM, Chapple C (2001) The *sng2* mutant of *Arabidopsis* is defective in the gene encoding the serine carboxypeptidase-like protein sinapoylglucose:choline sinapoyltransferase. Plant Journal 28: 83–94

Stehle F, Brandt W, Schmidt J, Milkowski C, Strack D (2008) Activities of Arabidopsis sinapoylglucose:malate sinapoyltransferase shed light on functional diversification of serine carboxypeptidase-like acyltransferases. Phytochemistry 69: 1826–1831

Stevenson PC, Nicolson SW, Wright GA (2017) Plant secondary metabolites in nectar: impacts on pollinators and ecological functions. Functional Ecology 31: 65–75

Velásquez AC, Chakravarthy S, Martin GB (2009) Virus-induced gene silencing (VIGS) in *Nicotiana benthamiana* and tomato. JoVE e1292

Whelan S, Goldman N (2001) A general empirical model of protein evolution derived from multiple protein families using a maximum-likelihood approach. Molecular Biology and Evolution 18: 691–699

